# Refactoring the upper sugar metabolism of *Pseudomonas putida* for co-utilization of disaccharides, pentoses, and hexoses

**DOI:** 10.1101/284182

**Authors:** Pavel Dvořák, Víctor de Lorenzo

**Author notes:** Corresponding author: Prof. V. de Lorenzo, Systems and Synthetic Biology Program, Centro Nacional de Biotecnología (CNB-CSIC), Darwin 3, Campus de Cantoblanco, Madrid 28049, Spain.

## Abstract

Given its capacity to tolerate stress, NAD(P)H/ NAD(P) balance, and increased ATP levels, the platform strain *Pseudomonas putida* EM42, a genome-edited derivative of the soil bacterium *P. putida* KT2440, can efficiently host a suite of harsh reactions of biotechnological interest. Because of the lifestyle of the original isolate, however, the nutritional repertoire of *P. putida* EM42 is centered largely on organic acids, aromatic compounds and some hexoses (glucose and fructose). To enlarge the biochemical network of *P. putida* EM42 to include disaccharides and pentoses, we implanted heterologous genetic modules for D-cellobiose and D-xylose metabolism into the enzymatic complement of this strain. Cellobiose was actively transported into the cells through the ABC complex formed by native proteins PP1015-PP1018. The knocked-in β-glucosidase BglC from *Thermobifida fusca* catalyzed intracellular cleavage of the disaccharide to D-glucose, which was then channelled to the default central metabolism. Xylose oxidation to the dead end product D-xylonate was prevented by by deleting the *gcd* gene that encodes the broad substrate range quinone-dependent glucose dehydrogenase. Intracellular intake was then engineered by expressing the *Escherichia coli* proton-coupled symporter XylE. The sugar was further metabolized by the products of *E. coli xylA* (xylose isomerase) and *xylB* (xylulokinase) towards the pentose phosphate pathway. The resulting *P. putida* strain co-utilized xylose with glucose or cellobiose to complete depletion of the sugars. These results not only show the broadening of the metabolic capacity of a soil bacterium towards new substrates, but also promote *P. putida* EM42 as a platform for plug-in of new biochemical pathways for utilization and valorization of carbohydrate mixtures from lignocellulose processing.

## 1. Introduction

Due to the physicochemical stresses that prevail in the niches in which the soil bacterium *Pseudomonas putida* thrives (it is typically abundant in sites contaminated by industrial pollutants), this microorganism is endowed with a large number of traits desirable in hosts of harsh biotransformations of industrial interest (Nikel et al., 2014). The *P. putida* strain KT2440 is a saprophytic, non-pathogenic, GRAS-certified (Generally Recognized as Safe) bacterium; as the most thoroughly characterized laboratory pseudomonad, it has an expanding catalogue of available systems and synthetic biology tools (Aparicio et al., 2017; Elmore et al., 2017; Martínez-García and de Lorenzo, 2017, p.; Nogales et al., 2017). This bacterium is becoming a laboratory workhorse as well as a valued cell factory (Benedetti et al., 2016; Loeschcke and Thies, 2015; Nikel and de Lorenzo, 2013; Poblete-Castro et al., 2012). Its high resistance to endogenous and exogenous insults makes it tolerant to industrially relevant chemicals (*e.g.,* ethanol, *p*-coumaric acid, toluene) and to by-products of biomass hydrolysis (furfural, 5-(hydroxymethyl)furfural, benzoate, acetic acid) at concentrations that are inhibitory to other microbial platforms, including *Escherichia coli* (Calero et al., 2017; Guarnieri et al., 2017; Johnson and Beckham, 2015; Nikel and de Lorenzo, 2014). The nutritional landscape of typical *P. putida* niches (plant rhizosphere, polluted soil) has pushed its metabolic specialization towards aromatic compounds (Jiménez et al., 2002) and organic acids (Dos Santos et al., 2004). The very few carbohydrates on which *P. putida* KT2440 can grow are confined to some hexoses (glucose and fructose), with an inability to metabolize disaccharides or 5-carbon sugars productively (Nogales et al., 2017; Puchałka et al., 2008; Rojo, 2010). This limits the options for its use as a platform to process the carbohydrate products of cellulosic and lignocellulosic waste.

Lignocellulose can be decomposed to cellulose (25-55%), hemicellulose (11-50%), and lignin (10-40%), all of which can be further hydrolyzed enzymatically to shorter carbohydrate polymers and oligomers, monomeric sugars, and lignin-derived aromatics (Mosier et al., 2005). When standard commercial cellulosic cocktails are applied, D-glucose and D-xylose are two major monomeric products of pretreated plant biomass hydrolysi (Taha et al., 2016). A number of microorganisms have been engineered to use these sugars as substrates for the biomanufacturing of value-added chemicals (Kawaguchi et al., 2016). Cellodextrins, including D-cellobiose, are the most abundant by-products of cellulose saccharification and predominate following partial hydrolysis (Chen, 2015; Singhania et al., 2013). Well defined, industrially relevant microbes with efficient cellobiose metabolism are thus very desirable (Kawaguchi et al., 2016; Parisutham et al., 2017; Taha et al., 2016). Both cellobiose and xylose metabolism have been established artificially in a number of microorganisms (Ha et al., 2011; Lane et al., 2015; Le Meur et al., 2012; Lee et al., 2016; Meijnen et al., 2008; Shin et al., 2014; Vinuselvi and Lee, 2011) and some *Pseudomonas* species can use xylose as a C source (Liu et al., 2015). Nonetheless, the native capacity of *P. putida* KT2440 to host transformations of lignin-derived aromatics (Linger et al., 2014) has not been combined with co-consumption of these major lignocellulose-derived sugars.

Here we sought to engineer *P. putida* for efficient growth on cellobiose and to test its ability to co-metabolize this disaccharide with xylose. We combined a metabolic engineering approach (**Fig. 1**) with the *P. putida* KT2440-derived strain *P. putida* EM42 (Martínez-García et al., 2014b). This strain has a streamlined genome (300 genes, ∼4.3 % of the genome deleted) that results in improved physiological properties compared to *P. putida* KT2440 including lower sensitivity to oxidative stress, increased growth rates, and enhanced expression of heterologous genes (Lieder et al., 2015; Martínez-García et al., 2014b). We show below that recruitment of one heterologous β-glucosidase for intracellular cellobiose hydrolysis, and the implementation of three enterobacterial genes (xylose transporter, isomerase and kinase) sufficed to cause disaccharide and the pentose co-utilization by *P. putida* EM42 with inactivated glucose dehydrogenase while maintaining its ability to use glucose. We also demonstrate that *P. putida* metabolism generates more ATP when cells are grown on cellobiose instead of glucose. This study expands the catalytic scope of *P. putida* towards utilization of major components of all three lignocellulose-derived fractions. Moreover, given that the cellobiose, as the bulk by-product of standard cellulose saccharification, frequently remains untouched in the sugar mix (due to the inability of most microorganisms to assimilate it), our results demonstrate rational tailoring of an industrially relevant microbial host to achieve a specific step in such a biotechnological value chain.

**Figure 1.**
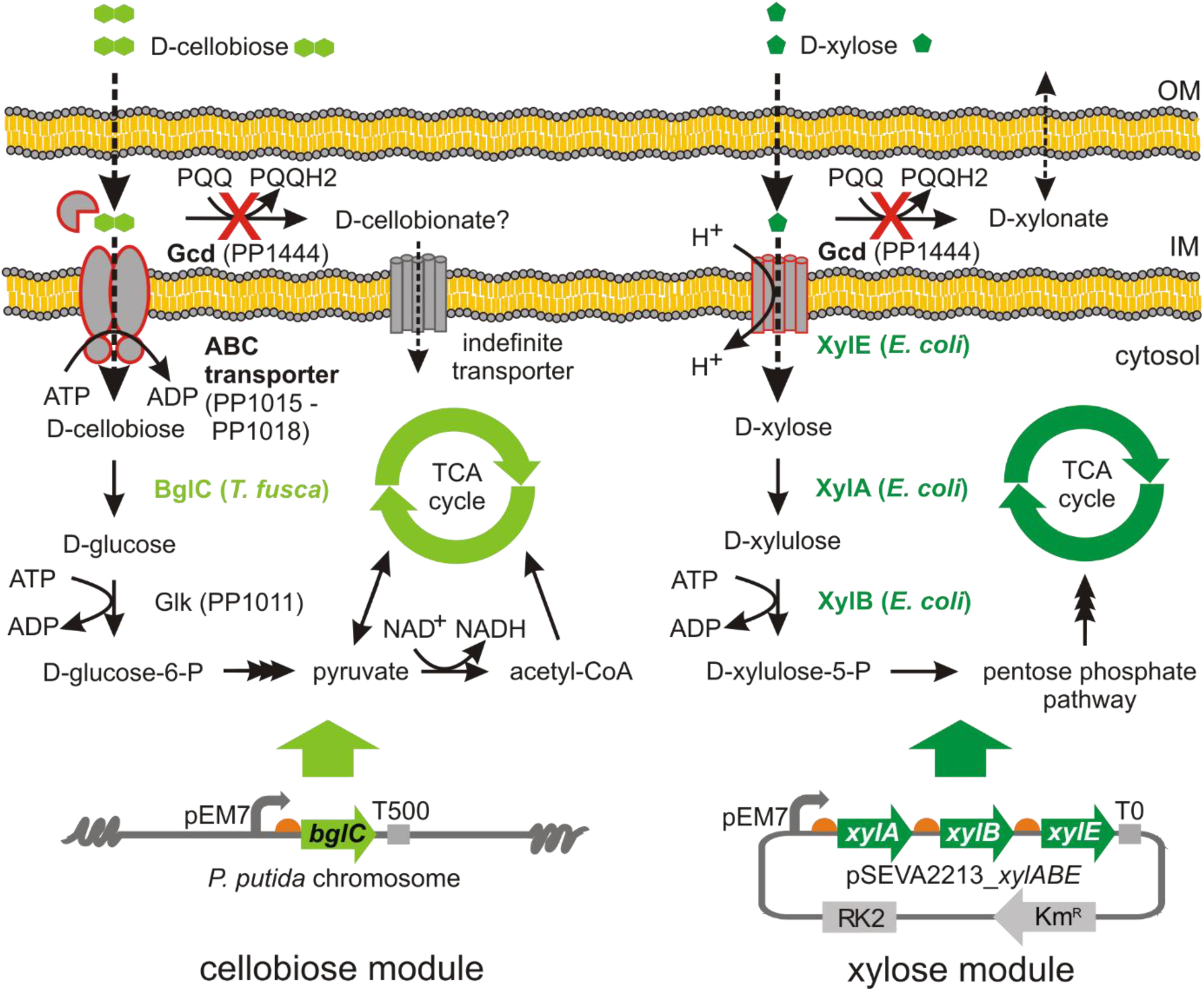
Engineering *Pseudomonas putida* EM42 for co-utilization of D-cellobiose and D-xylose. Cellobiose metabolism in *P. putida* was established by implantating the β-glucosidase BglC from *Thermobifida fusca*. Experiments suggested that cellobiose enters *P. putida* cells through the same pathways as glucose, (i) the ATP-dependent ABC transporter, and (ii) the peripheral oxidative route, which starts with periplasmic conversion of cellobiose to putative intermediate D-cellobionate through the action of the membrane-bound glucose dehydrogenase Gcd. Xylose metabolism in *P. putida* was established by implantating of xylose isomerase XylA, xylulokinase XylB, and xylose-proton symporter XylE from *E. coli*. For the purpose of cellobiose and xylose co-utilization, an expression cassette with the *bglC* gene was inserted into the chromosome of *P. putida* EM42 Δ*gcd*, while the synthetic *xylABE* operon was expressed from the low copy pSEVA2213 plasmid under the constitutive pEM7 promoter. The EM42 Δ*gcd* mutant was used to avoid xylose conversion by glucose dehydrogenase to dead-end product D-xylonate. PQQ, pyrroloquinoline quinone; Glk, glucokinase; TCA cycle, tricarboxylic acid cycle; OM, (outer membrane); IM (inner membrane).

## 2. Materials and methods

### 2.1. Bacterial strains, plasmids, and growth conditions

All bacterial strains and plasmids used in this study are listed in **Table 1**. *Escherichia coli* strains used for cloning or triparental mating were routinely grown in lysogeny broth (LB; 10 g L^-1^ tryptone, 5 g L^-1^ yeast extract, 5 g L^-1^ NaCl) with agitation (170 rpm) at 37°C. Cloramphenicol (Cm, 30 μg mL^-1^) was supplemented to the medium with *E. coli* helper strain HB101. *Pseudomonas putida* recombinants were routinely pre-cultured overnight in 2.5 mL of LB medium with agitation of 300 rpm (Heidolph Unimax 1010 and Heidolph Incubator 1000; Heidolph Instruments, Germany) at 30°C. For initial tests of expression of heterologous genes in *P. putida*, cells were transferred to 25 mL of fresh LB medium in Erlenmeyer flask and cultivated as described in section 2.4. For the growth experiments with different carbohydrates, overnight culture was spun by centrifugation (4,000 g, RT, 5 min), washed with M9 minimal medium (per 1 L: 4.25 g Na_2_HPO_4_ 2H_2_O, 1.5 g KH_2_PO_4_, 0.25 g NaCl, 0.5 g NH_4_Cl) added with MgSO_4_ to the final concentration of 2 mM, and with 2.5 mL L^-1^ trace element solution (Abril et al., 1989). Thiamine HCl (1 mM) was added to the minimal medium for cultures with *E. coli* recombinants. Cells were resuspended to OD_600_ of 0.1 in 25 mL of the same medium with kanamycin (Km, 50 µg mL^-1^), in case of recombinants with pSEVA2213 or pSEVA238 plasmid, or streptomycin (Sm, 60 µg mL^-1^), in case of *P. putida* EM42 Δ*gcd bglC*, and with carbon source (glucose, xylose, or cellobiose) of concentration defined in the text or respective figure caption. All used solid media (LB and M9) contained 15 g L^-1^ agar. M9 solid media were prepared with 2 mM MgSO_4_, 2.5 mL L^-1^ trace element solution (Abril et al., 1989) and 0.2 % (w/v) citrate, 0.4 % xylose, or 0.4 % cellobiose used as a sole carbon source.

**Table 1.**
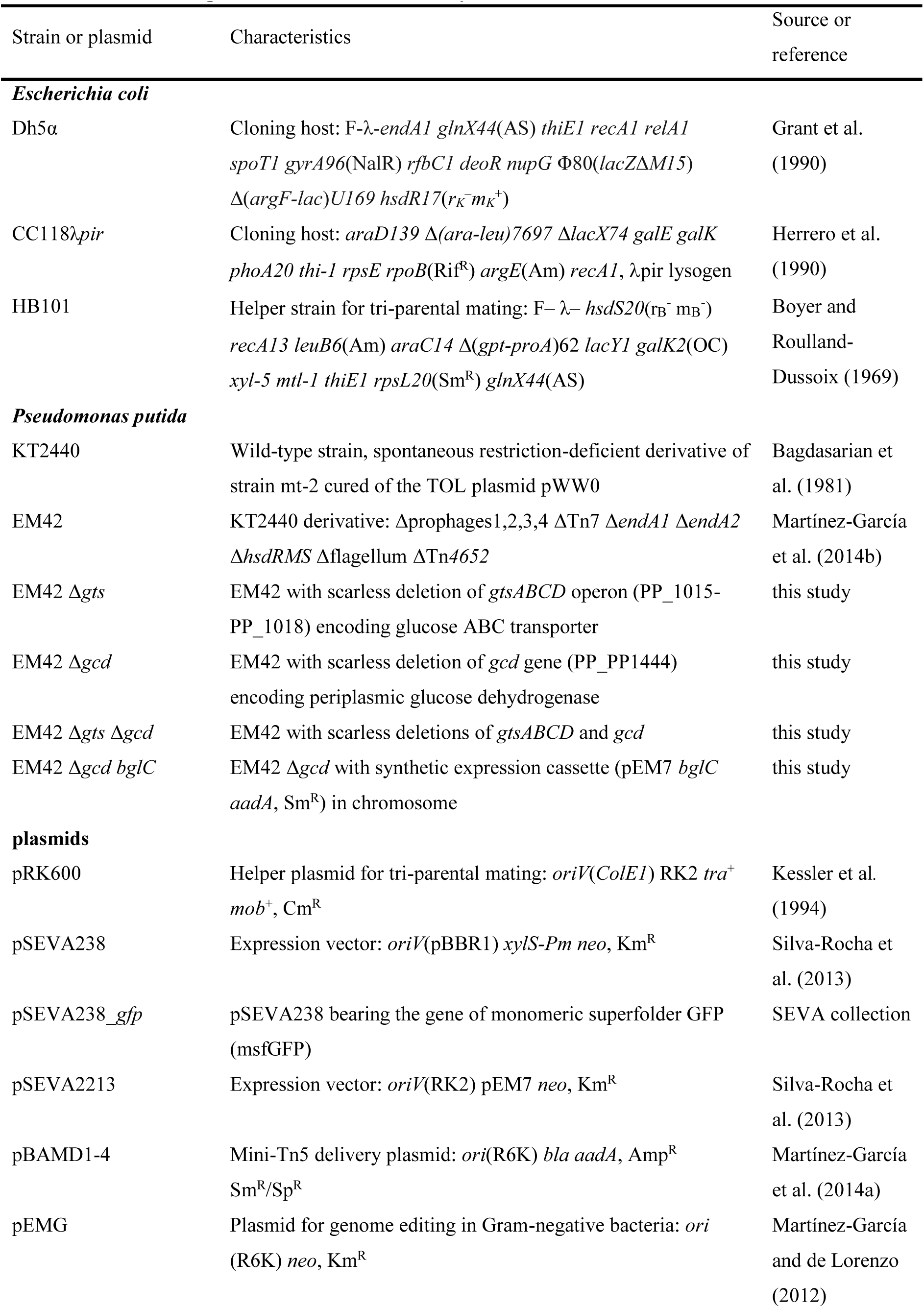
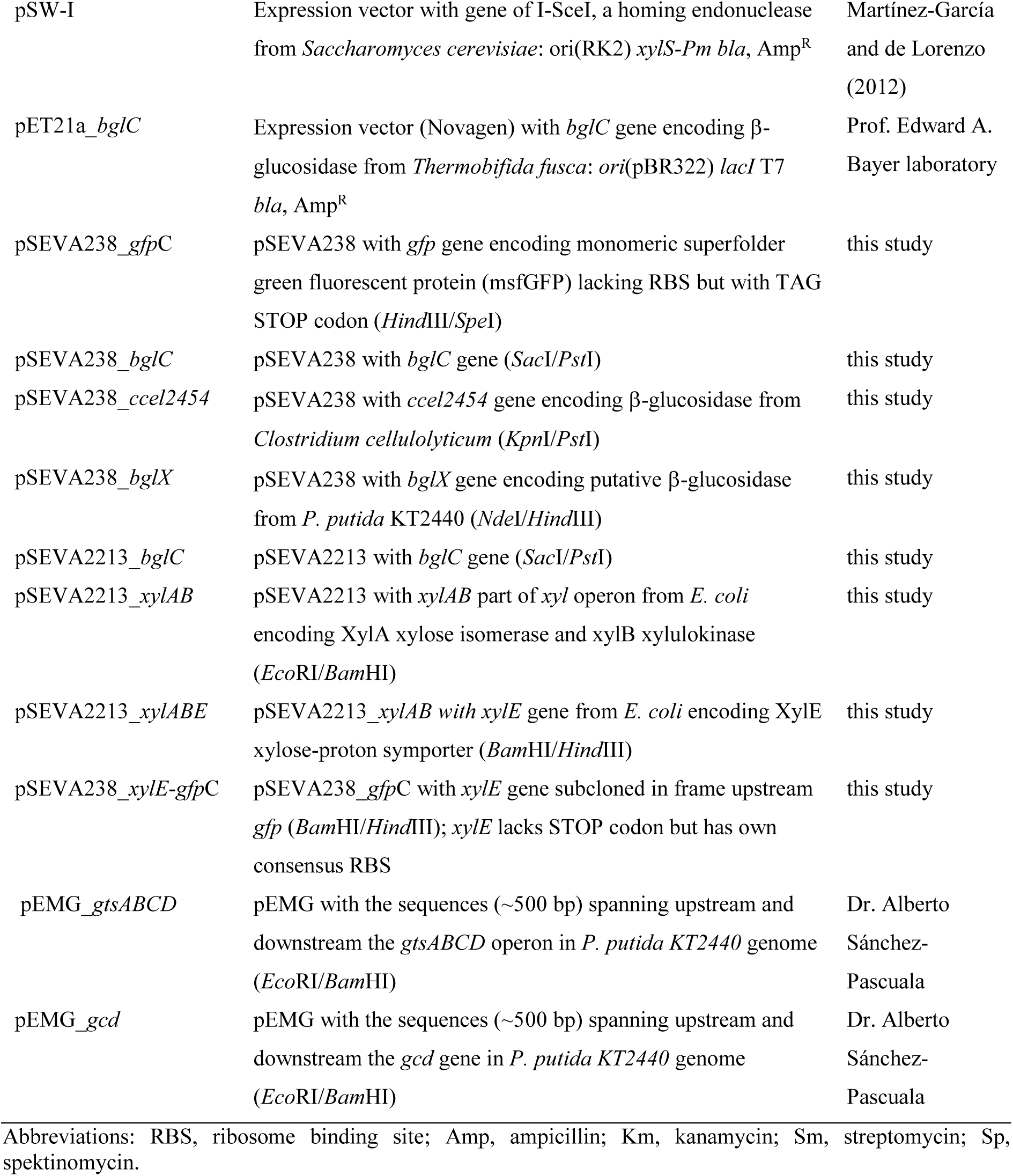
Strains and plasmids used in this study.

### 2.2. Plasmid and strain constructions

DNA was manipulated using standard laboratory protocols (Sambrook and Russell, 2001). Genomic DNA was isolated using GenElute bacterial genomic DNA kit (Sigma-Aldrich, USA). Plasmid DNA was isolated with QIAprep Spin Miniprep kit (Qiagen, USA). The oligonucleotide primers used in this study (**Table S1**) were purchased from Sigma-Aldrich (USA). The genes of interest were amplified by polymerase chain reaction (PCR) using Q5 high fidelity DNA polymerase (New England BioLabs, USA) according to the manufacturer’s protocol. The reaction mixture (50 μL) further contained polymerase HF or GC buffer (New England BioLabs, USA), dNTPs mix (0.2 mM each; Roche, Switzerland), respective primers (0.5 mM each), water, template DNA, and DMSO. GC buffer and DMSO were used for amplification of genes from *P. putida*. PCR products were purified with NucleoSpin Gel and PCR Clean-up (Macherey-Nagel, Germany). DNA concentration was measured with NanoVue spetrophotometer (GE Healthcare, USA). Colony PCR was performed using 2x PCR Master Mix solution of Taq DNA polymerase, dNTPs and reaction buffer (Promega, USA). All used restriction enzymes were from New England BioLabs (USA). Digested DNA fragments were ligated using Quick Ligation kit (New England BioLabs, USA). PCR products and digested plasmids separated by DNA electrophoresis with 0.8 % (w/v) agarose gels were visualised using Molecular Imager VersaDoc (Bio-Rad, USA). Plasmid constructs were confirmed by DNA sequencing (Macrogen, South Korea). Chemocompetent *E. coli* Dh5α cells were transformed with ligation mixtures or complete plasmids and individual clones selected on LB agar plates with Km (50 µg mL^-1^) were used for preparation of glycerol (20 % w/v) stocks. Constructed plasmids were transferred from *E. coli* Dh5α donor to *P. putida* EM42 by tripartite mating, using *E. coli* HB101 helper strain with pRK600 plasmid (**Table 1**). Alternatively, electroporation (2.5 kV, 4 – 5 ms pulse) was used for transformation of *P. putida* cells with selected plasmids using a MicroPulser electroporator and Gene Pulser Cuvettes with 0.2 cm gap (Bio-Rad, USA). Preparation of *P. putida* electrocompetent cells and electroporation procedure was performed as described elsewhere (Aparicio et al., 2015). *P. putida* transconjugants or transformants were selected on M9 agar plates with citrate or LB agar plates, respectively, with Km (50 µg mL^-1^) at 30°C overnight.

#### Construction of cellobiose metabolism module

The *ccel_2454* gene encoding β-glucosidase (EC 3.2.1.21) from *Clostridium cellulolyticum* was synthesized together with consensus ribosome binding site (RBS; GeneArt/Thermo Fisher Scientific, Germany) and subcloned from delivery vector pMA_*ccel2454* into pSEVA238 upon digestion with *Kpn*I and *Pst*I resulting in pSEVA238_*ccel2454*. The β-glucosidase encoding *bglX* gene was PCR amplified from the genomic DNA of *P. putida* KT2440 using bglX fw and bglX rv primers, digested with *Nde*I and *Hind*III and cloned into corresponding restriction sites of modified pSEVA238 resulting in pSEVA238_*bglX*. The *Nde*I site and a consensus RBS were previously introduced into the standard SEVA polylinker of pSEVA238 (unpublished plasmid). The *bglC* gene encoding β-glucosidase from *Thermobifida fusca* with N-terminal 6xHis tag was subcloned from pET21a_*bglC* construct into *Nde*I and *Hind*III restriction sites of modified pSEVA238. The *bglC* gene with the RBS and His tag was subsequently PCR amplified using primers bglC fw and bglC rv, the PCR product was cut with *Sac*I and *Pst*I and subcloned into pSEVA2213 giving rise to pSEVA2213_*bglC*.

#### Insertion of bglC gene into P. putida chromosome

The *bglC* gene with consensus RBS was subcloned into *Sac*I and *Pst*I sites of mini-Tn5-vector pBAMD1-4 (Martínez-García et al., 2014a). Original pBAMD1-4 plasmid was endowed with pEM7 promoter subcloned into *Avr*II and *Eco*RI sites. *P. putida* EM42 Δgcd cells (100 μL) were electroporated with plasmid DNA (200 ng) and recovered for 7 h in 5 mL of modified Terrific Broth (TB) medium (yeast extract 24 g L^-1^, tryptone 20 g L^-1^, KH_2_PO_4_ 0.017 M, K_2_HPO_4_ 0.072 M) at 30°C with shaking (170 rpm). Cells were collected by centrifugation (4000 rpm, 10 min) and resuspended in 100 mL of selection M9 medium with 5 g L^-1^ cellobiose and streptomycin (50 μg mL^-1^). After four days of incubation at 30°C with shaking (170 rpm), cells were spun (4000 rpm, 15 min) and plated on selection M9 agar plates with 5 g L^-1^ cellobiose and streptomycin (50 μg mL^-1^). Three fastest growing clones were re-streaked on fresh M9 agar plates with streptomycin or with streptomycin (50 μg mL^-1^) and ampicillin (500 µg mL^-1^) to rule out insertion of the whole pBAMD1-4 plasmid. The growth of three candidates in liquid minimal medium with cellobiose was verified. The insertion site of expression cassette (pEM7 promoter, *bglC* gene, T500 transcriptional terminator, and *aadA* gene) in chromosome of the fastest growing clone was determined by two-round arbitrary primed PCR with Arb6, Arb2, ME-O-Sm-Ext-F, and ME-O-Sm-Int-F primers (**Table S1**) following the protocol described before (Martínez-García et al., 2014a). Me-O-Sm-Int-F was used as a sequencing primer for PCR product. Position of *bglC* expression cassette in *P. putida* chromosome was reversly verified by colony PCR with bglC check fw and xerD check rv primers.

#### Construction of xylose metabolism module

The *xylAB* part of *E. coli xyl* operon encoding xylose isomerase (EC 5.3.1.5) XylA and xylulokinase (EC 2.7.1.17) XylB was amplified from genomic DNA of *E. coli* BL21 (DE3) using xylAB fw and xylAB rv primers. PCR product was digested with *Eco*RI and *Bam*HI and ligated into pSEVA2213 giving rise to pSEVA2213_*xylAB*. The gene of xylose-proton symporter (*xylE*) was amplified from the genomic DNA of *E.coli* BL21 (DE3) using two-step PCR protocol. In the first step, the gene was amplified using xylE fw 1 and xylE rv 1 primers. The sample of the reaction mixture with the PCR product (1 μL) was transferred into the second reaction with xylE fw 2 and xylE rv 2 primers. Final PCR product was digested with *Bam*HI and *Hind*III and cloned downstream *xylAB* operon in pSEVA2213_*xylAB* resulting in pSEVA2213_*xylABE*. For the purpose of construction of the plasmid allowing translational fusion of XylE to monomeric superfolded GFP (msfGFP), *gfp* gene was initially amplified without its own RBS but with STOP codon from pSEVA238_*gfp* plasmid (SEVA collection) using gfpC fw and gfpC rv primers. The PCR product was digested with *Hind*III and *Spe*I and ligated into pSEVA238, cut with the same pair of enzymes, giving rise to pSEVA238_gfpC. The *xylE* gene was amplified from pSEVA2213_*xylABE* with its synthetic RBS but without STOP codon using xylE-gfp fw and xylE-gfp rv primers. The PCR product was digested with *Bam*HI and *Hind*III and cloned upstream the *gfp* gene in pSEVA238_*gfp*C, resulting in pSEVA238_*xylE*-*gfp*C.

#### Preparation of deletion mutants of P. putida EM42

Deletion mutants were prepared using the protocol described previously (Aparicio et al., 2015). Briefly, the regions of approximately 500 bp upstream and downstream the *gtsABCD* genes (PP_1015 – PP_1018) were PCR amplified with TS1F-gtsABCD, TS1R-gtsABCD and TS2F-gtsABCD, TS2R-gtsABCD primers, respectively. TS1 and TS2 fragments were joined through SOEing-PCR (Horton et al., 1990), the PCR product was digested with *Eco*RI and *Bam*HI and cloned into non-replicative pEMG plasmid. The resulting pEMG_*gtsABCD* construct was propagated in *E. coli* CC118λ*pir* cells and the whole TS1-TS2 region was sequenced in several clones selected based on results of colony PCR with TS1F-gtsABCD and TS2R-gtsABCD primers (product of about 1 kb expected). The sequence verified pEMG_*gtsABCD* plasmid was transformed into competent EM42 cells by electroporation. Transformants were selected on LB agar plates with Km and co-integrates were identified by colony PCR with TS1F-gtsABCD and TS2R-gtsABCD primers. The pSW-I plasmid was transformed into selected co-integrate by electroporation. Transformants were plated on LB agar plates with Km and Amp and expression of I-SceI in selected clone inoculated into 5 mL of LB was induced with 1 mM 3-methylbenzoic acid (3MB) overnight. Induced cells were plated on LB agar plates with Amp and the positive clone EM42 Δ*gts* with a loss of Km resistance marker and deletion was confirmed by colony PCR using check(-)gtsABCD fw and TS2R-gtsABCD primers. PCR product size in case of scarless deletion was 1250 bp. The quinoprotein glucose dehydrogenase gene *gcd* (PP_1444) was deleted accordingly in *P. putida* EM42, resulting in *P. putida* EM42 Δ*gcd*, and in *P. putida* EM42 Δ*gts*, resulting in *P. putida* EM42 Δ*gts* Δ*gcd*, using a set of TS primers listed in **Table S1**. Expression of I-SceI in selected co-integrates was induced with 1 mM 3MB for 6 hrs. Check(-)gcd fw and check(-)gcd rv primers were used to confirm deletion of *gcd* gene. PCR product size in case of deletion was 1500 bp. *P. putida* recombinants were cured of pSW-I plasmid after several passes in LB medium lacking Amp.

### 2.3. Calculations of dry cell weight and growth parameters

Biomass was determined as dry cell weight. Samples of cultures grown in M9 minimal medium with 5 g L^-1^ glucose were transferred into 2 mL pre-dried and pre-weighed Eppendorf tubes and pelleted at 13,000 g for 10 min. The pellets were washed twice with distilled water and dried at 80°C for 48 h. Based on the prepared standard curve, one A_600_ unit is equivalent to 0.38 g L^-1^ of dry cell weight. Specific growth rate (μ) was determined during exponential growth as a slope of the data points obtained by plotting the natural logarithm of A_600_ values against time. Substrate consumption rate (*r*) was determined for initial 12 and 24 h of culture as *r* = (c substrate at *t*_0_ −c substrate at *t*_1_) / (*t*_1_ - *t*_0_). Biomass yield (Y_X/S_) was determined 24 h after each culture started to grow exponentially as Y_X/S_ = c biomass at *t*_1_ / (c substrate at *t*_0_ - c substrate at *t*_1_). Specific carbon (C) consumption rate (*q*_s_) was determined during exponential growth on glucose or cellobiose as *q*_s_ = (mmol C at *t*_0_ - mmol C at *t*_1_) / ((*t*_1_ - *t*_0_) * (g biomass at *t*_1_ - g biomass at *t*_0_)).

### 2.4. Enzyme activity assays

For initial screening of β-glucosidase activities, 25 mL of LB medium was inoculated from overnight cultures to A_600_ = 0.05 and cells were grown for 3 h at 30°C with shaking (170 rpm). Expression of β-glucosidase genes from pSEVA238 plasmid was then induced with 1 mM 3MB. After induction, cells were grown in the same conditions for additional 5 h and then harvested by centrifugation (4,000 g, 4°C, 10 min). Cell pellets were lysed by adding 1 mL of BugBuster Protein Exctraction Reagent with 1 µL of Lysonase Bioprocessing Reagent (both from Merck Millipore, USA) for 15 min at RT with slow agitation. For later β-glucosidase activity measurement of BglC in EM42 recombinants, cell lysates were prepared by spinning (21,000 g, 4°C, 2 min) 4 mL of cells growing in 25 mL of minimal medium with 5 g L^-1^ cellobiose (except for *P. putida* EM42 Δgts Δgcd pSEVA2213_*bglC* recombinant, which was grown in LB medium). Cells were collected in mid log phase (A_600_ = 1.0). Cell pellets were added with 200 uL of BugBuster Protein Exctraction Reagent and 0.2 µL of Lysonase Bioprocessing Reagent and lysed for 15 min at RT with slow agitation. Cell lysates for xylose isomerase and xylulokinase activity determination were prepared by sonicating concentrated cell solutions prepared by spinning (4,000 g, 4°C, 10 min) 25 mL of cells grown in LB medium to A_600_ = 1.0. Cell pellets were washed by 5 mL of ice-cold 50 mM Tris-Cl buffer (pH 7.5) resuspended in 1 mL of the same buffer, placed in ice bath and disrupted by sonication. In all cases, cell lysates were centrifuged at 21,000 g for 30 min at 4°C and supernatants, termed here as cell-free extracts (CFE), were used for activity determination. Total protein concentration in CFE was measured using the method of Bradford (Bradford, 1976) with a commercial kit (Sigma-Aldrich, USA). Crystalline bovine serum albumin (Sigma-Aldrich, USA) was used as a protein standard.

β-glucosidase activity was measured using synthetic substrate *p*-nitrophenyl-*β*-D-glucopyranoside (pNPG; Sigma-Aldrich, USA). Reaction mixture of total volume = 600 μL contained 550 μL of 50 mM sodium phosphate buffer (pH 7.0), 30 μL of pNPG (final conc. 5 mM), and 20 μL of CFE. Reaction was started by adding CFE to the mixture of buffer and substrate in Eppendorf tube pre-incubated 10 min at 37°C. CFE from *P. putida* cells producing BglC enzyme was diluted 100-200 times. Reaction was terminated after 15 min of incubation at 37°C in thermoblock by adding 400 μL of 1 M Na_2_CO_3_. Linearity of the enzymatic reaction during 15 min time course was initially verified by periodical withdrawal of the samples from reaction mixture of total volume of 1800 μL. Absorbance of the mixture was measured at 405 nm with UV/Vis spectrophotometer Ultrospec 2100 (Biochrom, UK) and activity was calculated using calibration curve prepared with p-nitrophenol standard (Sigma-Aldrich, USA). β-glucosidase activity in culture supernatants was measured correspondingly with 166 μL of culture supernatant in 600 μL of reaction mixture.

Activity of xylose isomerase (XylA) was measured as described by Le Meur and co-workers (2012) in microtiter plate format. In this assay, activity of XylA is coupled to consumption of NADH by sorbitol dehydrogenase. The assay mixture of total volume of 200 μL contained 50 mM Tris-Cl buffer (pH 7.5), 1 mM triethanolamine, 0.2 mM NADH, 0.5 U of sorbitol dehydrogenase, 10 mM MgSO_4_, and 50 mM xylose. Reaction at 30°C was started by addition of 5 μL of CFE.

Xylulokinase (XylB) activity was determined using the assay described by Eliasson *et al*. (2000) in microtiter plate format. In this assay, XylB activity is coupled with activities of pyruvate kinase and lactate dehydrogenase leading to the consumption of NADH. The reaction mixture of total volume of 200 μL contained 50 mM Tris-Cl buffer (pH 7.5), 0.2 mM NADH, 2 mM ATP, 2 mM MgCl_2_, 0.2 mM phosphoenolpyruvate, 10 U of pyruvate kinase and 10 U, lactate dehydrogenase, and 10 mM D-xylulose. Reaction at 30°C was started by addition of 5 μL of 20-fold diluted CFE.

Both in xylose isomerase and xylulokinase assay, the depletion of NADH was measured spectrophotometrically at 340 nm with Victor^2^ 1420 Multilabel Counter (Perkin Elmer, USA). Molar extinction coefficient of 6.22 mM^-1^ cm^-1^ for NADH was used for activity calculations. 1 unit (U) of activity corresponds to 1 μmol of substrate (pNPG or NADH) converted by enzyme per 1 min.

Activity of glucose dehydrogenase (Gcd) in *P. putida* EM42 and *P. putida* EM42 Δgcd was determined by measuring conversion of 5 g L^-1^ xylose to xylonate by cell suspension of A_600_ = 0.55 in 25 mL of M9 medium at 30°C. The time course of the reaction was 6 h. 1 U of enzyme activity corresponds to 1 μmol of xylonate produced per 1 minute.

### 2.5. SDS-PAGE and Western blot analyses

CFE for determination of expression levels of selected enzymes were prepared using cell pellets from cultures induced with 1 mM 3MB and lysed with BugBuster Protein Exctraction Reagent as described above. Samples of CFE containing 5 μg of total protein were added with 5x Laemmli buffer, boiled at 95°C for 5 min and separated by SDS-PAGE using 12 % gels. CFE prepared from *P. putida* cells with empty pSEVA238 plasmid was used as control. Gels were stained with Coomassie Brilliant Blue R-250 (Fluka/Sigma-Aldrich, Switzerland).

The staining step was omitted for Western blotting. Instead, proteins were electrotransferred from gel onto Immobilon-P membrane (Merck Millipore, Germany) of pore size = 0.45 μm using Trans-Blot SD Semi-Dry Transfer Cell (Bio-Rad, USA). Transfer conditions were: constant electric current of 0.1 A per gel, voltage of 5-7 V, time of run 30 min. Membrane was blocked overnight at 4°C in 3 % (w/v) dry milk in PBS buffer with 1 % (v/v) TWEEN 20 and then incubated with mouse anti-6xHis tag monoclonal antibody-HRP conjugate (Clontech, USA) for 2 h at RT. Membrane was washed with PBS buffer with 1 % (v/v) TWEEN 20 and the proteins were visualized after incubation with BM Chemiluminescence Blotting Substrate (POD; Roche, Switzerland) using Amersham Imager 600 (GE Healthcare Life Sciences, USA).

### 2.6. Determination of ATP levels in P. putida cells

The ATP content in *P. putida* recombinants growing in M9 minimal medium with 5 g L^-1^ glucose or cellobiose was determined as described previously by Lai *et al*. (2016). Briefly, 2 mL of cell culture of A_600_ = 0.5, representing ∼0.38 mg CDW, was centrifuged (13,000 g, 4°C 2 min). Pellets were resuspended in 250 µL of ice-cold 20 mM Tris-Cl buffer (pH 7.75) with 2 mM EDTA and ATP was extracted using the trichloroacetic acid (TCA) method (Lundin and Thore, 1975). Ice-cold 5 % (w/v) TCA (250 μL) with 4 mM EDTA was added, the suspension was mixed by vortexing for 20 s and incubated on ice for 20 min. Then, the suspension was centrifuged (13,000 g, 4°C, 10 min) and supernatant (10 μL) was diluted 20-fold with ice cold 20 mM Tris-Cl buffer (pH 7.75) with 2 mM EDTA. The solution (50 μL) was mixed with 50 μL of the reagent from bioluminescence based ATP determination kit (Biaffin, Germany) prepared according to the manufacturer’s instructions. After 10 min of incubation in dark, the luminescence signal of diluted supernatants was read in white 96-well assay plate (Corning Incorporated, USA) using the microplate reader SpectraMax (Molecular Devices, USA). The luminescence was quantified using the calibration curve prepared with pure ATP (Sigma-Aldrich, USA).

### 2.7. Confocal microscopy

Localization of XylE transporter fused with msfGFP in the cell membrane was verified by confocal microscopy of cells expressing the chimeric gene from pSEVA238_*xylE*-*gfp*C. Cells bearing pSEVA238_*gfp* or empty pSEVA238 were used as controls. Cells were grown in LB medium (30°C, 275 rpm) until A_600_ = 0.5. Expression was induced with 0.5 mM 3MB and the growth continued at the same conditions for another 2.5 h. The sample of cell culture (100 μL) was centrifuged (5,000 g, 4°C, 5 min). Cells were washed twice with 1.5 mL of ice cold phosphate buffer saline (PBS; per 1 L: 8 g NaCl, 0.2 g KCl, 1.44 g Na_2_HPO_4_, 0.24 g KH_2_PO_4_, pH adjusted to 7.4 with HCl) and finally resuspended in 1 mL of the same buffer. Cells (5 μL of suspension) were mounted on poly-L-lysine coated glass slides (Sigma-Aldrich, USA) for 20 min, covered with cover glass and the slides were analyzed using confocal multispectral miscroscope Leica TCS SP5 (Leica Microsystems, Germany).

### 2.8. Other analytical techniques

Cell growth was monitored by measuring absorbance of a cell suspension at 600 nm using UV/Vis spectrophotometer Ultrospec 2100 (Biochrom, UK). For determination of glucose, xylose, xylonate, and cellobiose concentrations, culture supernatant (0.5 mL) was centrifuged (21,000 g, 4°C, 10 min), filtered using Whatman Puradisc 4 mm syringe filter with nylon membrane (GE Healthcare Life Sciences, USA) and stored at −20°C for following analyses. Samples were analyzed by HPLC-LS system 920LC with light scattering (PL-ELS) detector (Agilent Technologies, USA) equipped with Microsorb MV NH2 column (5 µm, 250 mm x 4.6 mm). MilliQ H_2_O (A) and acetonitrile (B) were used as eluents at a flow rate of 1 mL min^-1^. Column temperature was 30°C. Chemicals were identified using pure compound standards. Glucose and xylose concentrations in culture supernatants were also determined by Glucose (GO) Assay Kit (Sigma-Aldrich, USA) and Xylose Assay Kit (Megazyme, Ireland), respectively. Xylonic acid was measured using the hydroxamate method (Lien, 1959). Samples of culture supernatants (75 μL) were mixed 1:1 with 1.3 M HCl and heated at 100°C for 20 min to convert xylonate to xylono-γ-lactone. Samples were cooled down on ice and 50 μL were added to 100 μL of hydroxylamine reagent freshly prepared by mixing 2 M hydroxylamine HCl with 2 M NaOH (pH of the reagent should be between 7 and 8). After 2 min interval, 65 μL of 3.2 M HCl and subsequently 50 μL of FeCl_3_ solution (10 g in 100 mL of 0.1 M HCl) were added. Absorbance at 550 nm was measured immediately using Victor^2^ 1420 Multilabel Counter (Perkin Elmer, USA). Xylonate concentrations were quantified with a standard curve prepared using the pure compound (Sigma-Aldrich, USA).

### 2.9. Statistical analyses

All the experiments reported here were repeated independently at least twice (number of repetitions is specified in figure and table legends). The mean values and corresponding standard deviations (SD) are presented. When appropriate, data were treated with a two-tailed Student’s t test in Microsoft Office Excel 2013 (Microsoft Corp., USA) and confidence intervals were calculated for given parameter to manifest a statistically significant difference in means between two experimental datasets.

## 3. Results and discussion

### 3.1. Engineering the cellobiose metabolism module

Initial tests with *P. putida* EM42 in minimal medium with cellobiose showed no growth of this platform strain on the disaccharide (**Fig. 2A**). The absence of cellobiose assimilation implied either lack of transport through the cell membranes or missing enzymatic machinery for disaccharide hydrolysis or phosphorolysis. As we detected no β-glucosidase (EC 3.2.1.21) activity in culture supernatants or lysates prepared from cells incubated with cellobiose, we focused initially on the latter bottleneck. Previous attempts to engineer bacteria for cellobiose utilization and valorization included periplasmic or extracellular β-glucosidase expression (Chen et al., 2011; Rutter et al., 2013), or its surface display (Muñoz-Gutiérrez et al., 2014). This last approach was also recently used for *P. putida* KT2440, but surface display of three cellulases, including β-glucosidase BglA from *Clostridium thermocellum*, resulted in only trace conversion of cellulosic substrate into glucose (Tozakidis et al., 2016). Alternative strategies included intracellular assimilation of cellobiose *via* endogenous or exogenous β-glucosidase (Ha et al., 2011; Vinuselvi and Lee, 2011), and *via* phosphorolysis catalyzed by cellobiose phosphorylase (Shin et al., 2014). Despite the need for an efficient cellobiose transporter, intracellular assimilation is generally regarded as more efficient, because it circumvents carbon catabolite repression (CCR) and prevents cellobiose inhibition of extracellular cellulases in the bioreactor (Ha et al., 2011; Parisutham et al., 2017; Teugjas and Väljamäe, 2013).

**Figure 2.**
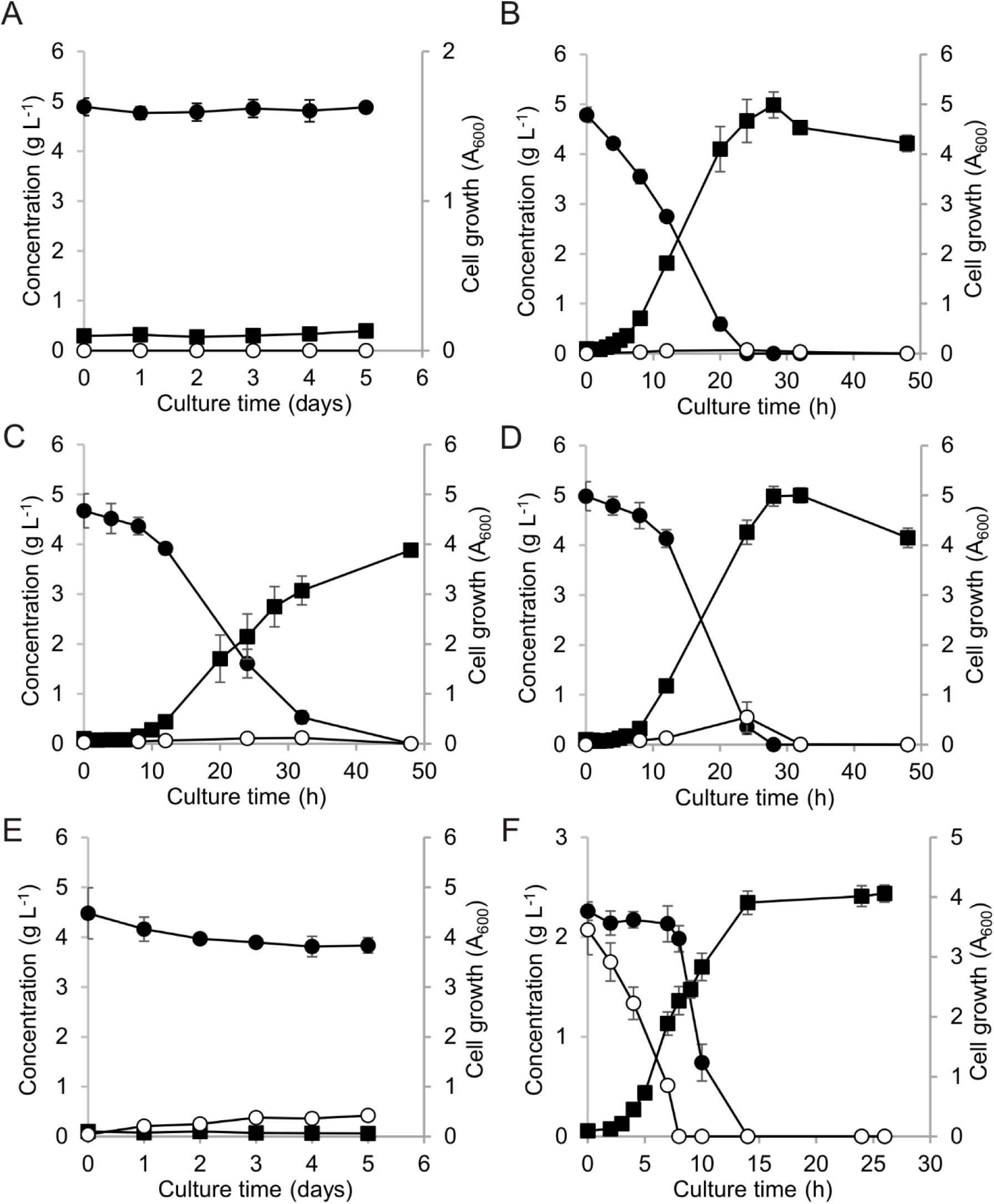
Growth of *P. putida* EM42 and its recombinants in minimal medium with 5 g L^-1^ D-cellobiose. Experiments were carried out in shaken flasks (30°C, 170 rpm). (A) *P. putida* EM42, (B) *P. putida* EM42 pSEVA2213_*bglC*, (C) *P. putida* EM42 Δ*gts* pSEVA2213_*bglC*, (D) *P. putida* EM42 Δ*gcd* pSEVA2213_*bglC*, (E) *P. putida* EM42 Δ*gts* Δ*gcd* pSEVA2213_*bglC*, (F) *P. putida* EM42 pSEVA2213_*bglC* in minimal medium with D-glucose and D-cellobiose (2 g L^-1^ each). D-cellobiose, filled circles (●); D-glucose, open circles (○); cell growth, filled squares (▪).Data points shown as mean ± SD from two to three independent experiments.

In *P. putida* EM42, we probed functional expression of two different intracellular β-glucosidases, Ccel_2454 from the Gram-positive mesophilic bacterium *Clostridium cellulolyticum* and BglC from the Gram-positive thermophilic bacterium *Thermobifida fusca* (see **Supporting information** for nucleotide sequences of all enzymes used in this study). Both enzymes have been expressed successfully in *E. coli* and are compatible with moderate temperatures and neutral or slightly acidic pH (Desai et al., 2014; Fan et al., 2012; Spiridonov and Wilson, 2001). In addition, we overexpressed the EM42 endogenous *bglX* gene (PP_1403), which is annotated as periplasmic β-glucosidase in the Pseudomonas Genome Database (www.pseudomonas.com/). The enzyme has relatively high amino acid sequence identity (61 %) with the well-characterized *E. coli* β-glucosidase BglX (Yang et al., 1996). Each of these genes was cloned into the pSEVA238 plasmid downstream of the inducible XylS/Pm promoter. We tested the effect of gene expression on EM42 viability, as well as soluble protein production and enzyme activity in cell-free extracts (CFE).

All three enzymes were produced in the soluble fraction of the *P. putida* chassis grown in LB medium (**Fig. S1A**), but only Ccel_2454 and BglC showed measurable β-glucosidase activity. No activity was detected in CFE containing endogenous BglX, whose overexpression also had a clear toxic effect on the host (**Fig. S2**). Absence of BglX β-glucosidase activity might be explained by loss of protein function due to a mutation(s) gained during a period when the bacterium was not benefited by the enzyme. The Ccel_2454 activity (0.03 ± 0.01 U/1 mg of total protein in CFE) measured with the colorimetric substrate p-nitrophenol-beta-D-glucopyranoside at 37°C (pH 7.0) was limited. In contrast, CFE of *P. putida* EM42 expressing the *bglC* gene showed high activity (4.57 ± 0.50 U mg^-1^). Trace BglC activity was also detected in culture supernatants (2.76 ± 0.24 U L^-1^), which indicates that a small amount of overexpressed enzyme can exit the cell during growth through an unknown mechanism. Western blot analysis of a CFE sample containing His-tagged BglC and a sample of culture supernatant (**Fig. S1B**) nonetheless confirmed that the great majority of the protein was expressed intracellularly. After induction with 0.2 mM 3-methylbenzoate (3MB), strong BglC expression and activity enabled growth of the host strain in minimal medium with 5 g L^-1^ cellobiose as a sole carbon source. In this preliminary shake flask experiment, the OD_600_ at 48 h was 3.6 (data not shown).

The *bglC* gene was subsequently subcloned into the low copy plasmid pSEVA2213 with the constitutive promoter pEM7 that functions well in both *P. putida* and *E. coli* (Zobel et al., 2015). The *P. putida* EM42 bearing the pSEVA2213_*bglC* construct grew rapidly in minimal medium with 5 g L^-1^ cellobiose, with only a ∼2 h adaptation period and a specific growth rate of 0.35 ± 0.02 h^-1^ (**Fig. 2B**, **Table 2**). The substrate consumption rate and biomass yield parameters were about 40 % and 20 % lower, respectively, than those of *P. putida* EM42 pSEVA2213 cultured in minimal medium with glucose (**Table 2**). As all disaccharide was consumed within 24 h of culture, the *P. putida* strain outperformed the best engineered *E. coli* strain CP12CHBASC30, which assimilated 3.3 g L^-1^ of cellobiose after 32 h in comparable conditions (Vinuselvi and Lee, 2011). No growth was observed of *E. coli* Dh5α transformed with pSEVA2213_*bglC* and incubated in the same conditions, despite the relatively high β-glucosidases activity (1.51 ± 0.12 U mg^-1^) detected in these cells. This result highlights the excellent match in between the BglC cellulase selected and the *P. putida* host.

**Table 2.**
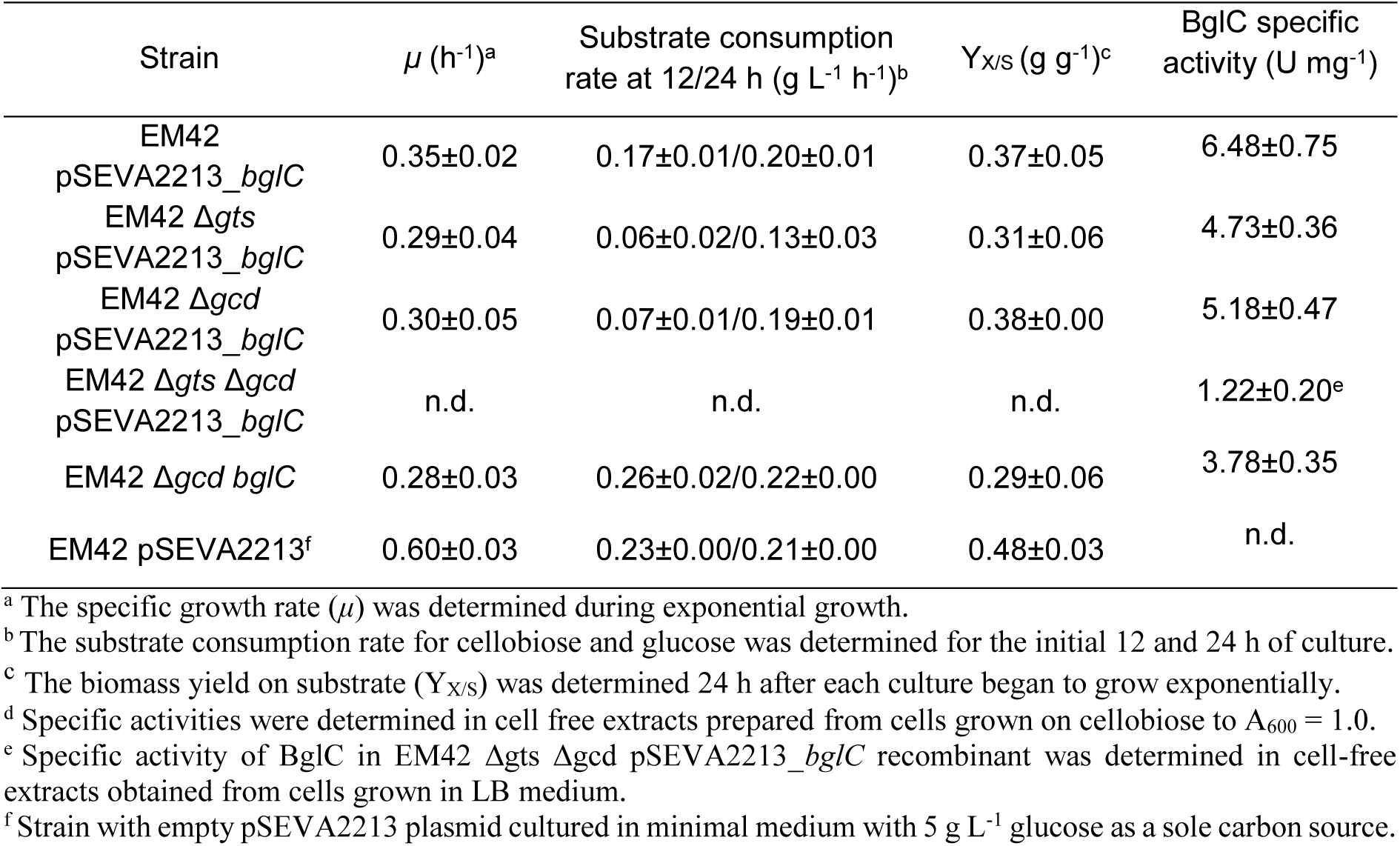
Growth parameters for batch cultures of *Pseudomonas putida* EM42 recombinants carried out on cellobiose. Values shown as mean ± SD from two to three independent experiments.

### 3.2. Understanding cellobiose transport in P. putida EM42

To decipher transport pathways for cellobiose (a glucose dimer) in *P. putida*, we first focused on the well-described importation of monomeric glucose (del Castillo et al., 2007). There are two routes for glucose assimilation in *P. putida* KT2440. The first route encompasses direct translocation of a glucose molecule from periplasm to cytoplasm by the ATP-dependent ABC transporter, encoded by the *gtsABCD* operon (PP_1015-PP_1018), and subsequent phosphorylation of hexose to glucose-6-phosphate by glucokinase (del Castillo et al., 2007). The second is a periplasmic pathway formed by membrane-bound PQQ-dependent glucose dehydrogenase Gcd (PP_1444). This enzyme oxidizes D-glucose to D-glucono-1,5-lactone, which is hydrolyzed either spontaneously or by the action of gluconolactonase Gnl (PP_1170) to D-gluconate. Gluconate can be further oxidized to 2-keto-D-gluconate by periplasmic gluconate dehydrogenase (Gad). Both gluconate and 2-ketogluconate can pass through the outer membrane or are imported into the cytoplasm, where the oxidation pathway merges with the direct phosphorylation route at the level of 6-phospho-D-gluconate (del Castillo et al., 2007).

Uptake of cellobiose and other cellodextrins of varying lengths through the ABC-type transporters is common in cellulolytic bacteria due to the relatively broad specificity of these systems (Nataf et al., 2009; Parisutham et al., 2017). To test its relevance for cellobiose uptake in EM42 strain, we deleted the *gtsABCD* operon that encodes the ABC glucose transporter. The mutant transformed with the pSEVA2213_*bglC* plasmid showed a substantially prolonged (∼7 h) adaptation phase on cellobiose (**Fig. 2C**). Three determined growth parameters were reduced when compared with *P. putida* EM42 pSEVA2213_*bglC* (**Table 2**). The results suggested that the glucose ABC transporter plays an important role in cellobiose uptake, but is not the only access route for the disaccharide in *P. putida*.

Closure of the second glucose uptake route by deleting the glucose dehydrogenase gene *gcd* in *P. putida* EM42 had no notable effect on growth on cellobiose, but slightly prolonged the lag phase (∼3 h; **Fig. 2D**). Substrate consumption during the initial 12 h of growth was nonetheless significantly reduced (**Table 2**), which might be attributed to the uptake of cellobiose only through the direct phosphorylation route and slower initial expression of ABC transporter components that is normally induced by monomeric glucose (del Castillo et al., 2007). It is worth noting here that neither the genes which encode two out of three carbohydrate-selective porins (*oprB-1* and *oprB-2*) adjacent to *gtsABCD* and *gcd*, respectively, nor their regulatory sequences were affected by the scarless deletions.

Growth on disaccharide was completely abolished when the deletions in the direct phosphorylation and oxidative routes were combined (**Fig. 2E**, **Table 2**). It can be thus argued that the peripheral glucose pathway also takes part in cellobiose assimilation by *P. putida*. One could speculate that Gcd also has cellobiose dehydrogenase activity (EC 1.1.99.18) and converts cellobiose to cellobiono-1,5-lactone (Henriksson et al., 2000). Much like glucono-1,5-lactone, in the presence of water cellobiono-1,5-lactone might be hydrolyzed spontaneously to cellobionic acid, which would be transported to the cytoplasm. Intracellular β-glucosidase could then cleave cellobionate to glucose and gluconic acid (Li et al., 2015), two molecules easily metabolized by *P. putida*. Nonetheless, neither cellobionic acid formation nor its further metabolism in *P. putida* can be confirmed based on currently available experimental data. Hence, the detailed functioning of the peripheral oxidative route in upper cellobiose metabolism in *P. putida* remains to be elucidated by our future experiments.

From the acquired growth parameters of these *P. putida* mutants, it can be deduced that the direct phosphorylation route is of major importance for cellobiose assimilation. On the other hand, experimental evidence shows that glucose enters *P. putida* cells predominantly through the peripheral oxidative pathway (Nikel et al., 2015). Despite these opposing access route preferences, when *P. putida* EM42 pSEVA2213_*bglC* was exposed to a mixture of the two sugars (2 g L^-1^ each), we observed diauxic growth (**Fig. 2F**). Glucose was utilized first during the initial 8 h of the experiment. When all hexose was removed from the medium, cellobiose was consumed rapidly during the next 6 h of the culture. To conclude, these experiments suggest that glucose and cellobiose share the same acess routes in *P. putida* and that monomeric hexose is a preferred substrate in the mixture of the two carbon sources.

### 3.3. Probing energetic benefit of cellobiose metabolism in engineered P. putida

ATP is a universal energy source and a major driving force for biochemical processes in microbial cell factories (Hara and Kondo, 2015). Due to its variant of glycolysis – the Entner-Doudoroff pathway - *P. putida* yields only one net ATP per one mole of assimilated glucose (Nikel et al., 2015). In the case, for instance, of *E. coli* with its characteristic Embden– Meyerhof–Parnas pathway, the ATP yield per molecule of glucose is twice as high. It is thought that environmental or engineered microorganisms that prefer to metabolize cellobiose instead of glucose are more energetic and robust than their glucose-utilizing counterparts (Chen, 2015; Lynd et al., 2002; Parisutham et al., 2017). The benefits were demonstrated in bacteria with a cellobiose-specific PEP-phosphotransferase system transporter, in which one mole of ATP is consumed per one mole of imported substrate; this was also apparent in microbes that metabolize cellobiose through phosphorolysis, in which only one ATP per disaccharide is needed to form two activated molecules of glucose (Kajikawa and Masaki, 1999; Shin et al., 2014; Thurston et al., 1993).

We determined ATP levels in EM42 pSEVA2213 and EM42 pSEVA2213_*bglC* strains grown on glucose and cellobiose, respectively, to evaluate the effect of altered substrate on *P. putida* energy status (**Fig. 3A**). Indeed, the ATP level in *P. putida* grown on cellobiose was almost double that of the cells cultured on glucose. ATP savings could partially stem from cellobiose transport. One ATP per molecule of glucose is theoretically saved when cellobiose enters the cell with the help of an ABC-type transporter, as the ATP cost is known to be constant per import event (Parisutham et al., 2017). The oxidative pathway is also thought to provide the cell with additional energy through the transfer of electrons from membrane-bound Gcd and Gad directly to the respiratory chain enzymes (Ebert et al., 2011). The higher ATP level on cellobiose was not accompanied by more efficient conversion of carbon into biomass (**Table 2**, **Fig. 3B**), which might be attributed to the higher respiration activity of the cells grown on cellobiose (Ebert et al., 2011). An extraordinary amount of ATP in the cell could also inhibit citrate synthase, which would slow down the TCA cycle as well as formation of biomass precursors (Smith and Williamson, 1971). To sum up, the ATP saved during the growth on a biotechnologically relevant substrate will further increase the value of the engineered EM42 platform strain as a cell factory for bioproduction.

**Figure 3.**
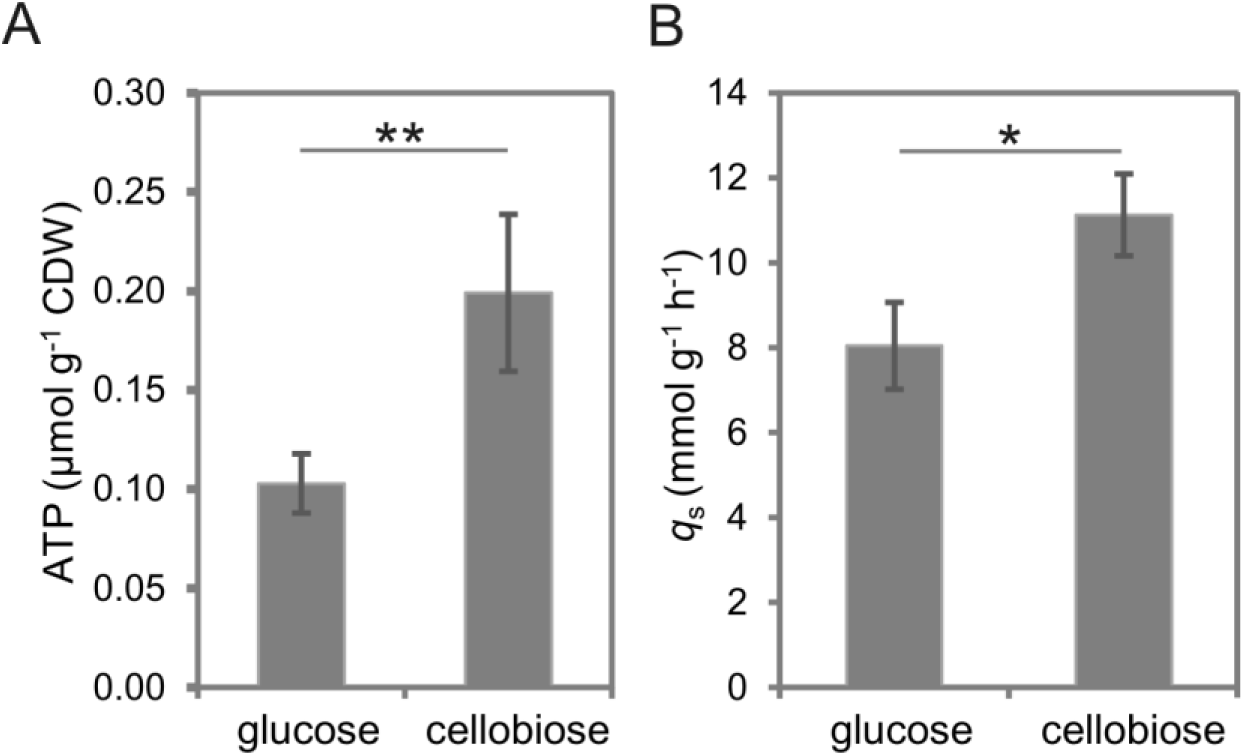
Energetic benefit of *P. putida* EM42 cells growing on D-cellobiose. (A) Comparison of intracellular ATP concentrations in *P. putida* EM42 pSEVA2213 growing on 5g L^-1^ D-glucose and in *P. putida* EM42 pSEVA2213_*bglC* growing in minimal medium with 5 g L^-1^ D-cellobiose. Measurements were performed with cells collected from cultures in exponential growth (A_600_ = 0.5). (B) Specific carbon consumption rate (*q*_s_) for glucose (8.04 ± 1.02) and cellobiose (11.13 ± 0.96) determined in the exponential growth phase for *P. putida* EM42 pSEVA2213 and *P. putida* EM42 pSEVA2213_*bglC*, respectively. Data shown as mean ± SD from three independent experiments. Asterisks denote significance in difference in between two means at P < 0.05 (*) or P < 0.01 (**).

### 3.4. Engineering metabolic module for xylose utilization

To test the capacity of *P. putida* to co-utilize cellobiose with pentoses, we established a D-xylose metabolism in our platform strain EM42. *Pseudomonas putida* KT2440 is unable to utilize xylose as a sole carbon source for growth (Le Meur et al., 2012; Nogales et al., 2017; Puchałka et al., 2008). Both KT2440 and *P. putida* S12, another pseudomonad with biotechnological potential, were nonetheless engineered successfully for xylose utilization by implementation of the isomerase pathway from *E. coli* (Le Meur et al., 2012; Meijnen et al., 2008). This route fuels the pentose phosphate pathway *via* the action of xylose isomerase (EC 5.3.1.5) and xylulokinase (EC 2.7.1.17), which convert D-xylose to D-xylulose-5-phosphate (**Fig. 1**). We aimed at following the same strategy in our study.

We first verified the absence of xylose catabolism in *P. putida* EM42. Our bioinformatic analysis confirmed that the KT2440 genome has no genes that encode homologues of *E. coli* XylA or XylB proteins. The EM42 strain was then incubated in minimal medium with xylose, and the cell density and substrate concentration were measured for five consecutive days (**Fig. 4A**). No growth was detected, despite the fact that only 10% of the starting xylose concentration was detected in the culture medium after five days. Meijnen and co-workers (2008) described a similar phenomenon for engineered *P. putida* S12. In that case, the majority of D-xylose was oxidized to the dead-end product D-xylonate by the periplasmic glucose dehydrogenase Gcd. In fact, xylonate concentrations determined in samples from the five-day experiment with the strain EM42 suggested that all xylose was converted to the acid, which was not assimilated by the cells (**Fig. 4A**). Xylonate formation was accompanied by a decrease in the culture pH from 7.0 to 6.2. This initial experiment provided additional evidence of Gcd broad substrate specificity in *P. putida* KT2440 and its derivatives. The phenomenon of xylonate formation was not discussed in the study by Le Meur and colleagues (2012), who implanted the isomerase pathway in the KT2440 strain.

**Figure 4.**
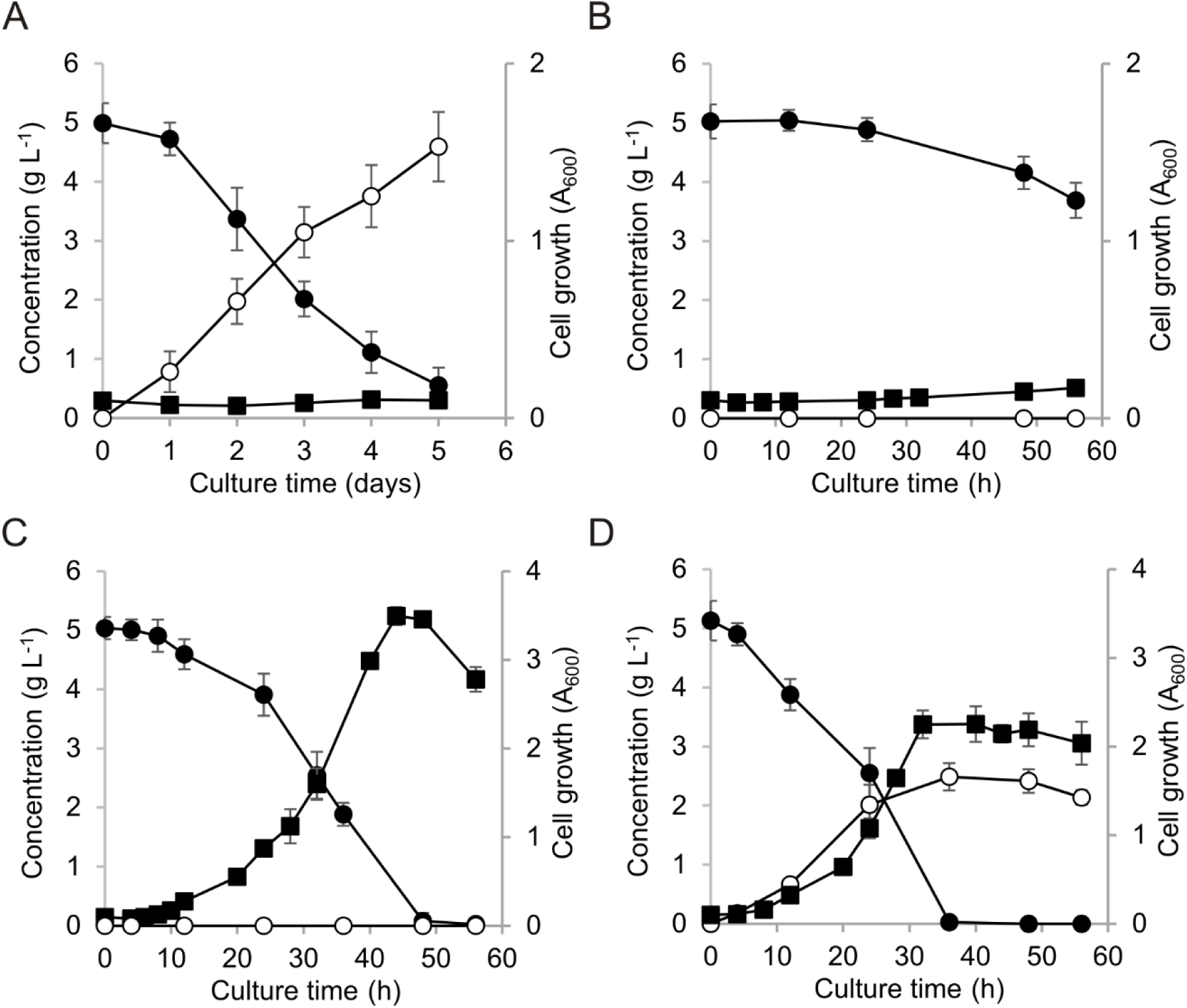
Growth of *P. putida* EM42 and its recombinants in minimal medium with 5 g L^-1^ D-xylose. Experiments were carried out in shaken flasks at 30°C and 170 rpm. (A) *P. putida* EM42, (B) *P. putida* EM42 Δ*gcd* pSEVA2213_*xylAB*, (C) *P. putida* EM42 Δ*gcd* pSEVA2213_*xylABE*, (D) *P. putida* EM42 pSEVA2213_*xylABE*. D-xylose, closed circles (●); D-xylonate, open circles (○); cell growth, closed squares (▪). Data shown as mean ± SD from three independent experiments.

To avoid accumulation of an undesirable metabolite, we transplanted the *xylAB* fragment of the *xyl* operon from *E. coli* BL21 (DE3), which encodes xylose isomerase XylA and xylulokinase XylB, directly to *P. putida* EM42 Δ*gcd*. The *xylAB* fragment was amplified as a whole. The *xylA* gene (**SI sequences**) was provided with a consensus RBS, with the native RBS maintained upstream of the *xylB* gene. The fragment was cloned into pSEVA2213 and *xylAB* expression was verified in the EM42 Δ*gcd* strain (**Fig. S3**). XylA and XylB activities determined in CFE were higher than those reported for engineered *P. putida* S12 growing on xylose (**Table 3**) (Meijnen et al., 2008). The recombinant EM42 cells nonetheless showed only limited growth and substrate uptake in minimal medium with 5 g L^-1^ xylose (**Table 3**, **Fig. 4B**). When xylonate accumulation no longer hindered efficient cell use of xylose, we found substrate transport to be another bottleneck to xylose metabolism in this host. This was not anticipated based on a previous study with the xylose-utilizing KT2440 strain, which reported only implantation of the XylAB metabolic module with no transport system (Le Meur et al., 2012). In another report, an upregulated glucose ABC transporter was nonetheless defined as one of the major changes that shaped laboratory-evolved *P. putida* S12 towards rapid growth on xylose (Meijnen et al., 2012).

**Table 3.** Growth parameters and specific activities of xylose isomerase and xylulokinase for batch cultures of *Pseudomonas putida* EM42 recombinants carried out on xylose. Data shown as mean ± SD from three independent experiments.

| Strain | $\mu$ (h<sup>-1</sup>)<sup>a</sup> | Substrate consumption rate (g L<sup>-1</sup> h<sup>-1</sup>)<sup>b</sup> | Y<sub>X/S</sub> (g g<sup>-1</sup>)<sup>c</sup> | XylA specific activity (U mg<sup>-1</sup>)<sup>d</sup> | XylB specific activity (U mg<sup>-1</sup>)<sup>d</sup> |
| --- | --- | --- | --- | --- | --- |
| EM42 Δ*gcd*<br>pSEVA2213_*xylAB* | 0.02±0.00 | 0.01±0.01 | 0.08±0.02 | 0.12±0.00 | 2.58±0.12 |
| EM42 Δ*gcd*<br>pSEVA2213_*xylABE* | 0.17±0.02 | 0.05±0.01 | 0.27±0.08 | 0.12±0.01 | 1.14±0.12 |

<sup>a</sup> The specific growth rate ( $\mu$ ) was determined during exponential growth.
<sup>b</sup> The xylose consumption rate was determined for the initial 24 h of culture.
<sup>c</sup> The biomass yield on substrate (Y<sub>X/S</sub>) was determined 24 h after each culture began exponential growth.
<sup>d</sup> Specific activities were determined in cell free extracts prepared from cells grown on xylose to OD<sub>600</sub> = 1.0.

The xylose-proton symporter XylE from *E. coli* is a relatively small (491 amino acids) single-gene transporter with a known tertiary structure and a well-described transport mechanism (Davis and Henderson, 1987; Wisedchaisri et al., 2014). In the recent study by Yim and coworkers (2016), XylE was selected as the best candidate among three tested pentose transporters that allowed growth of *Corynebacterium glutamicum* on xylose. The same transporter was also applied successfully in *Zymomonas mobilis* (Dunn and Rao, 2014). We probed XylE performance in *P. putida*. The gene was PCR-amplified from the genome of *E. coli* BL21(DE3) and supplied with consensus RBS to secure sufficient expression of *xylE* cloned downstream of the *xylAB* fragment in pSEVA2213_*xylAB*, to form the synthetic *xylABE* operon. The *xylE* was also simultaneously amplified without a stop codon and was cloned upstream of the *gfp* gene in the pSEVA238_*gfp* plasmid bearing the inducible XylS/Pm promoter to form translational fusion. The resulting construct was electroporated into *P. putida* EM42, and fluorescence microscopy confirmed targeting of XylE-GFP protein chimera in the cell membrane after 3MB induction (**Fig. 5**).

**Figure 5.**
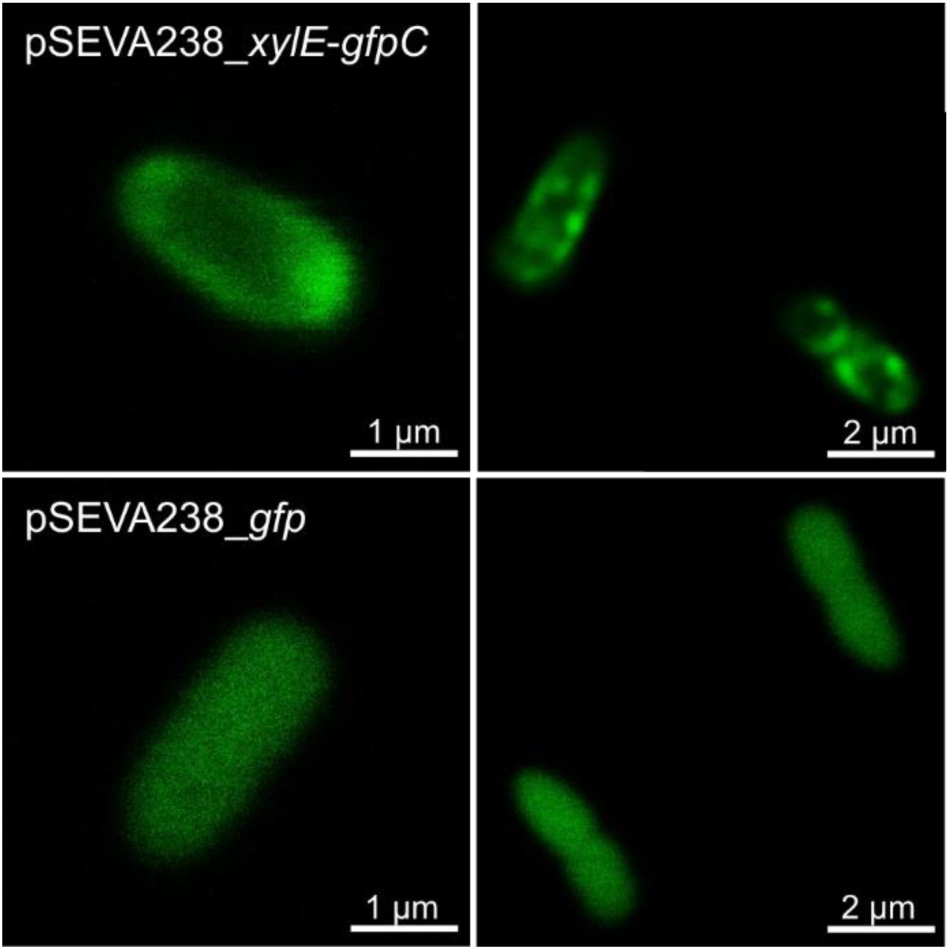
Localization of the XylE-GFP chimera in the cell membrane of *P. putida* EM42. The experiment was performed with *P. putida* EM42 pSEVA238_*xylE-gfpC* cells with *xylE* and *gfp* genes cloned to form a translational fusion, and with *P. putida* EM42 pSEVA238_*gfp* (negative control). Expression of the chimeric gene or *gfp* only was induced for 2.5 h with 0.5 mM 3-methylbenzoate in lysogeny broth at 30°C. GFP fluorescence in cells was then monitored with a confocal microscope. Note that GFP fluorescence in cells producing the XylE-GFP chimera is predominant in membrane regions.

Once we confirmed correct XylE expression and localization in *P. putida*, the EM42 Δ*gcd* strain was transformed with the pSEVA2213_*xylABE* construct to test functioning of the whole synthetic operon. Co-expression of the exogenous transporter with xylose isomerase and xylulokinase genes improved the specific growth rate on xylose by 8-fold when compared with the recombinant without *xylE* (**Fig. 4C** **and** **Table 2**); improvement was also observed for other growth parameters (**Table 2**). The faster growth of the recombinant bearing the *xylABE* synthetic operon could not be attributed to changes in XylA or XylB activitity (**Table 2**). We observed a >2-fold decrease in XylB activity when *xylE* was subcloned downstream of *xylB* and the position of the gene in operon was changed. Finally, we verified the importance of the *gcd* deletion for complete xylose utilization by *P. putida* recombinants in an experiment with the EM42 pSEVA2213_*xylABE* strain (**Fig. 4D**). It is clear from the time course of the culture that ∼50 % of all uptaken xylose was still converted non-productively to xylonate in the bacterium with functional Gcd, despite the presence of the heterologous machinery that funnels the substrate to the pentose phosphate pathway.

We thus demonstrate that efficient xylose metabolism can be established in the platform strain *P. putida* EM42 when two major bottlenecks – the peripheral oxidative pathway and the missing transport system – are removed. Neither of these bottlenecks was rationally engineered in previous studies of pseudomonad metabolization of xylose (Le Meur et al., 2012; Meijnen et al., 2008, 2012). The specific growth and xylose consumption rates of the *xylABE*-bearing recombinant were lower than those we measured for *P. putida* EM42 utilizing glucose or cellobiose. Both parameters can nonetheless be improved in co-utilization experiments in which host cell growth is supported by an additional carbon source (Ha et al., 2011; Lee et al., 2016).

### 3.5. Co-utilization of xylose with glucose and cellobiose by engineered P. putida EM42

Simultaneous uptake of carbohydrates is a desirable property in any microbial cell factory used in bioprocesses for valorization of lignocellulosic substrates (Lynd et al., 2002; Stephanopoulos, 2007). Co-utilization of biomass hydrolysis products increases the efficiency of the process and prevents accumulation of non-preferred sugars, usually pentoses, in batch and continuous fermentations (Jarmander et al., 2014; Kim et al., 2015). In industrially relevant microorganisms such as *E. coli*, *S. cerevisiae* or *Z. mobilis*, however, co-utilization of lignocellulose-derived sugars is hindered by complex CCR mechanisms that prioritize glucose from other carbon sources (Görke and Stülke, 2008; Kayikci and Nielsen, 2015). These mechanisms must be circumvented by mutagenesis or introduction of heterologous catabolic routes and sugar transporters (Kim et al., 2015; Lawford and Rousseau, 2002). As shown recently in yeast and *E. coli*, engineering microbes towards the use of cellobiose or other cellodextrins is a powerful alternative strategy for managing CCR (Ha et al., 2011; Vinuselvi and Lee, 2012). Glucose metabolism in *P. putida* is not as central as it is in *E. coli*, and in fact other substrates such as organic acids or amino acids are preferred to sugars (Rojo, 2010). Moreover, the isomerase pathway and the xylose transporter introduced into *P. putida* EM42 are of exogenous origin and their expression is not governed by the host. We thus anticipated that our recombinant strains would have co-utilized xylose with glucose or cellobiose with no restrictions.

We first sought to verify simultaneous utilization of glucose and xylose in *P. putida* EM42 Δ*gcd* pSEVA2213_*xylABE*. This experiment with two monomeric sugars was an essential prerequisite for co-utilization of xylose and cellobiose in engineered *P. putida*. The strain with the *gcd* deletion was used to avoid xylose oxidation to the dead-end by-product xylonate. As explained in section 3.2., this deletion is not detrimental either for glucose or for cellobiose uptake in *P. putida*, and both molecules can enter the cell with the help of ABC transporter. Equal concentrations of monosacharides (2 g L^-1^) were used to better visualize the differences in glucose and xylose consumption. *Pseudomonas putida* EM42 Δ*gcd* pSEVA2213_*xylABE* assimilated glucose and xylose simultaneously, and no sugar was detected in culture supernatants after 24 h (**Fig. 6B**). In cultures of the negative control *P. putida* EM42 Δ*gcd* pSEVA2213 lacking the *xylABE* operon, the xylose concentration dropped by only 13% in the same time period (**Fig. 6A**). It is possible that some xylose entered the cells by non-specific transport routes. The presence of the additional carbon source significantly accelerated xylose assimilation by recombinant *P. putida* (**Fig. 6B**). On average, 2 g L^-1^ of xylose were consumed during the initial 24 h of co-utilization experiments, while <1 g L^-1^ was mineralized in cultures with pentose alone at a starting concentration of 5 g L^-1^ (**Fig. 4C**). The substrate consumption rate of xylose was nonetheless still lower than that of glucose, which caused two-phase growth of the EM42 Δ*gcd* pSEVA2213_*xylABE* strain on two sugars at the same starting concentration (**Fig. 6B**). This experiment demonstrated the ability of engineered *P. putida* to co-utilize hexose and pentose without CCR.

**Figure 6.**
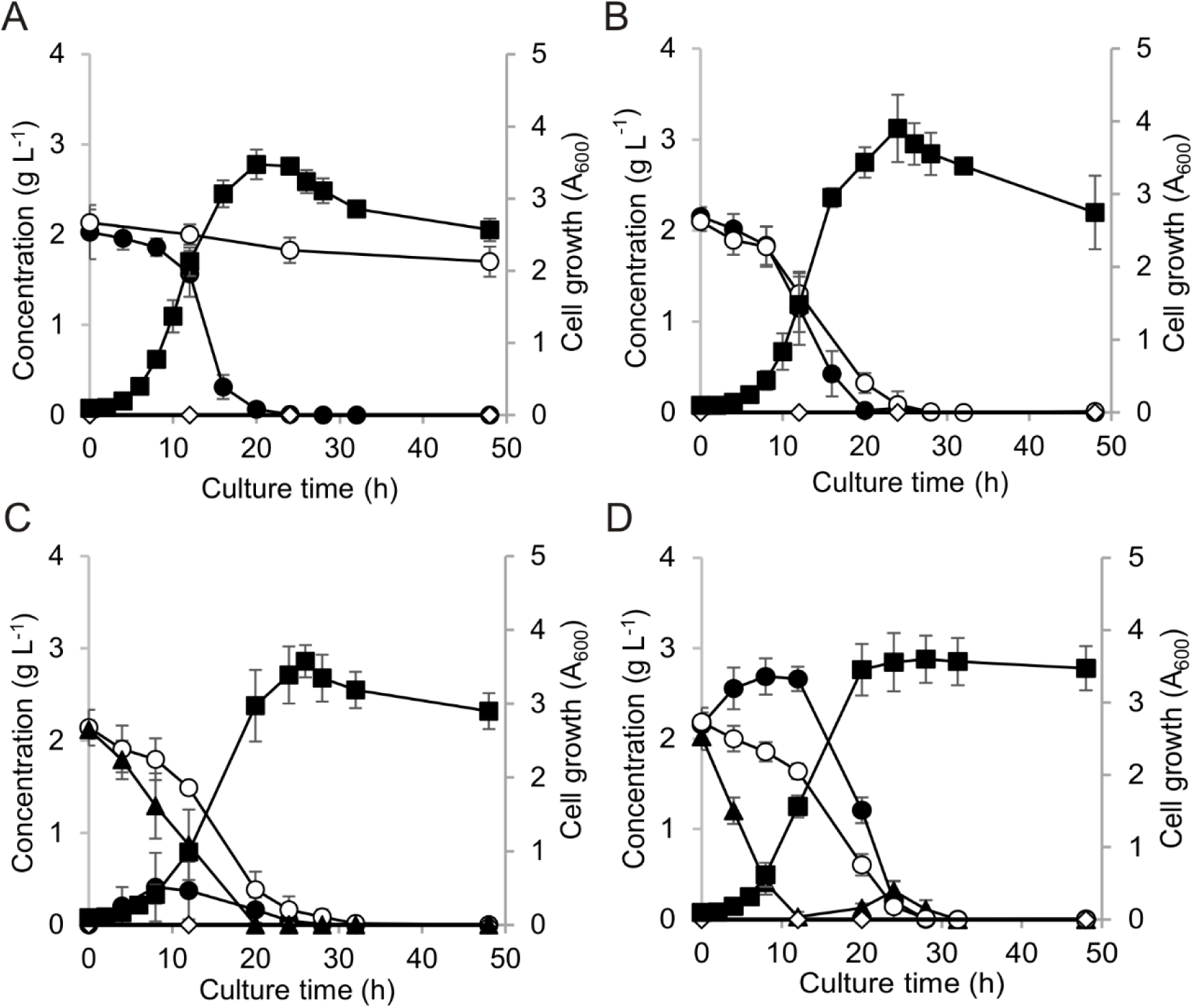
Co-utilization of D-xylose with D-glucose or D-cellobiose by *P. putida* EM42 recombinants in minimal medium. Experiments were carried out in shaken flasks (30°C, 170 rpm). (A) *P. putida* EM42 Δ*gcd* pSEVA2213 with glucose and xylose (2 g L^-1^ each), (B) *P. putida* EM42 Δ*gcd* pSEVA2213_*xylABE* with glucose and xylose (2 g L^-1^ each), (C) *P. putida* EM42 Δ*gcd bglC* pSEVA2213_*xylABE* with cellobiose and xylose (2 g L^-1^ each), (D) *P. putida* EM42 Δ*gcd bglC* pSEVA2213_*xylABE* with cellobiose, xylose, and glucose (2 g L^-1^ each). D-glucose, filled circles (●); D-xylose, open circles (○); D-cellobiose, filled triangles (▲); D-xylonate, open diamonds (◊); cell growth, filled squares (▪). Data shown as mean ± SD from three independent experiments.

For the cellobiose and xylose co-utilization experiments, new metabolic modules had to be combined in a single *P. putida* cell. As the synthetic *bglC*-*xylABE* operon borne on a single pSEVA2213 plasmid appeared to be unstable, we integrated the *bglC* gene directly into the *P. putida* EM42 Δ*gcd* chromosome. The gene with the pEM7 promoter and consensus RBS was subcloned into the mini-Tn5-vector pBAMD1-4, which allows for random chromosomal insertions and subsequent selection of the optimal phenotype from a broad expression landscape (Martínez-García et al., 2014a). The plasmid construct was electroporated into *P. putida* with a transformation efficiency of ∼160,000 CFU μg^-1^ of DNA. The best candidates were selected in minimal medium with cellobiose (see **Methods**). The fastest growing clone with the *bglC* insertion in gene annotated as the tyrosine recombinase subunit *xerD* (PP_1468) was designated *P. putida* EM42 Δ*gcd bglC*. With another tyrosine recombinase XerC, functional XerD is pivotal for the process of chromosome segregation during bacterial cell division (Blakely et al., 2000). Insertion of the expression cassette with *bglC* into the *xerD* gene nonetheless did not notably affect host fitness. *Pseudomonas putida* EM42 Δ*gcd bglC* growth parameters on cellobiose were comparable with those of the *P. putida* EM42 Δ*gcd* recombinant bearing the *bglC* gene on the plasmid; substrate consumption rate was even faster (**Table 2, Fig. S4**). The insertion had no effect on cell viability either in rich LB medium or in minimal medium with citrate as the gluconeogenic carbon source (**Fig. S5**).

Hence, *P. putida* EM42 Δ*gcd bglC* was transformed with the pSEVA2213_*xylABE* construct, and the resulting recombinant used for co-utilization experiments. Cells were incubated in minimal medium with cellobiose and xylose at equal concentrations (2 g L^-1^). Once again, both substrates were co-utilized rapidly; no residual carbohydrate was detected in supernatants after 28 h culture (**Fig. 6C**). Xylose was assimilated slower than cellobiose. We also tested performance of EM42 Δ*gcd bglC* pSEVA2213_*xylABE* strain in mixture of cellobiose, xylose, and glucose at 2 g L^-1^ concentration each (**Fig. 6D**). While xylose consumption remained unafected by presence of glucose in medium, cellobiose uptake was notably accelerated. In contrast, glucose concentration in the culture supernatants increased untill cellobiose was consumed and then all the hexose was co-utilized with remaining xylose. We argue that glucose, present in the medium already at the beginning of the culture, allowed faster cellobiose utilization by enhancing expression of the ABC transporter operon and adjacent *oprB* porin gene (del Castillo et al., 2007). We described in section 3.2. that the direct phosporylation route appears to be the major uptake pathway for cellobiose, but up to 80-90% of glucose is assimilated through the oxidative route in *P. putida* KT2440 (Nikel et al., 2015). Such induced expression of the ABC transporter in *P. putida* Δ*gcd* mutant could thus lead to the inverted substrate preference observed in the last co-utilization experiment.

Small amounts of extracellular glucose were detected in all the cultures with *P. putida* EM42 Δ*gcd* mutant grown on cellobiose (**Figs. 2D, 6C, 6D, and S4**). Cellobiose streamed into the cells only through the ABC transporter might cause temporal accumulation of intracellular glucose, which is released into the medium and later transported back to the cytoplasm. In contrast to many other β-glucosidases, BglC is not inhibited by glucose (Spiridonov and Wilson, 2001). We thus hypothesize that glucose accumulation in the EM42 Δ*gcd* mutant stems from an imbalance between β-glucosidase activity and *P. putida* glycolysis, namely its the upper part encompassing glucokinase (PP_1011) and glucose-6-phosphate 1-dehydrogenase (PP_1022, PP_4042, PP_5351). The system must accomodate all the glucose in its native form rather than its oxidized intermediates gluconate and 2-ketogluconate, which prevail in the cells with functional Gcd (Nikel et al., 2015). The imbalance could be reduced for instance by parallel modulation of expression of respective genes (Zhu et al., 2017).

## 4. Conclusions

Here we engineered *Pseudomonas putida* EM42, a robust platform strain derived from *P. putida* KT2440, to metabolize cellobiose and xylose, and to co-utilize these two carbohydrates. Within 24 h of culture in minimal medium, *P. putida* EM42 expressing the intracellular β-glucosidase BglC utilized 5 g L^-1^ cellobiose as a sole carbon source, thus outperforming the best cellobiose-utilizing *E. coli* strain constructed to date (Vinuselvi and Lee, 2011). This result highlights the need to select cellulases with good target host compatibility. We demonstrated that *P. putida* uses its native transport routes for cellobiose uptake, as no heterologous transporter had to be implanted into our recombinants. This aligns *P. putida* KT2440 and its derivatives with several other microorganisms, such as *Clostridium thermocellum*, *Klebsiella oxytoca*, *Neurospora crassa*, and *Streptomyces spp*., which possess cellodextrin transport systems and can thus manage cellobiose-rich mixtures that result from partial hydrolysis of cellulosic materials (Lynd et al., 2002; Nataf et al., 2009; Zhou et al., 2001). Metabolism of cellobiose generates more ATP in *P. putida* than when the same cells are cultured on glucose. The cellobiose-utilizing *P. putida* would thus be an even more robust host than the template strain for accommodating heterologous or engineering endogenous anabolic pathways for biosynthesis of value-added chemicals directly from the disaccharide or a co-substrate (Hara and Kondo, 2015). Finally, we identify the ability of *P. putida*, following introduction of the *xylABE* synthetic operon, to co-utilize cellobiose or glucose with pentose, with no need for further interventions in the regulatory mechanisms of central carbon metabolism. Xylose metabolism in the EM42 chassis was established based on the conclusions of previous reports of pentose-utilizing *P. putida* strains (Le Meur et al., 2012a; Meijnen et al., 2008a), whereas the need for an exogenous transporter and *gcd* deletion for complete xylose assimilation and co-utilization with glucose in our *P. putida* KT2440 derivative is demonstrated in this study.

Although there is indeed room for improvement and further testing of the strains constructed here, we argue that this study increases the value of *P. putida* for the biotechnological recycling of lignocellulosic feedstocks, specifically for processes that include partial hydrolysis of the input material. Using synthetic and systems biology approaches, carbon from new (hemic)cellulosic substrates-cellobiose and xylose-can be streamlined towards valuable chemicals whose production has been reported in *P. putida*, such as mcl-PHA (Poblete-Castro et al., 2013, p.), rhamnolipids (Tiso et al., 2017), terpenoids (Mi et al., 2014), coronatines (Gemperlein et al., 2017) and others (Loeschcke and Thies, 2015; Poblete-Castro et al., 2012). Given recent progress in *P. putida* KT2440 engineering for valorization of lignin (Johnson and Beckham, 2015; Linger et al., 2014), one could hypothesize that this effort could result in recombinant bacterial workhorses capable of simultaneous biotechnological processing of and adding value to all three lignocellulose-derived fractions.

## Acknowledgements

We would like to thank prof. Edward A. Bayer for providing us with pET21a_*bglC* plasmid, to Dr. Esteban Martínez-García for *P. putida* EM42 strain, and to Dr. Alberto Sánchez-Pascuala for pEMG_*gtsABCD* and pEMG_*gcd* plasmids and corresponding oligonucleotide primers. The project has received funding from the EU’s Horizon 2020 research and innovation programme under the Marie Sklodowska-Curie grant agreement No 704410 (FUTURE).

## Appendix A. Supplementary material

Dataset to this manuscript is available at https://data.mendeley.com/datasets with DOI: 10.17632/j7ypmvfnvt.1

