## Supplementary Materials for "Refactoring the upper sugar metabolism of *Pseudomonas putida* for co-utilization of disaccharides, pentoses, and hexoses"

* Corresponding author:

***Supplementary Figures***

**Figure S1.** Expression analysis of β-glucosidases tested in *Pseudomonas putida* EM42.

**Figure S2.** Effect of over-expression of tested β-glucosidases on growth of *P. putida* EM42 host.

**Figure S3.** Sodium dodecyl sulfate polyacrylamide gel electrophoresis of cell free extracts obtained from *P. putida* EM42 pSEVA2213 and *P. putida* EM42 pSEVA2213_*xylAB*.

**Figure S4.** Growth of *P. putida* EM42 Δ*gcd bglC* in minimal medium with 5 g $L^{-1}$ D-cellobiose.

**Figure S5.** Comparison of growth of *P. putida* EM42 Δ*gcd* and *P. putida* EM42 Δ*gcd bglC* in rich growth medium and in minimal medium with gluconeogenic carbon source.

***Supplementary Tables***

**Table S1.** Oligonucleotide primers used in this study.

***Nucleotide sequences of genes used in this study***

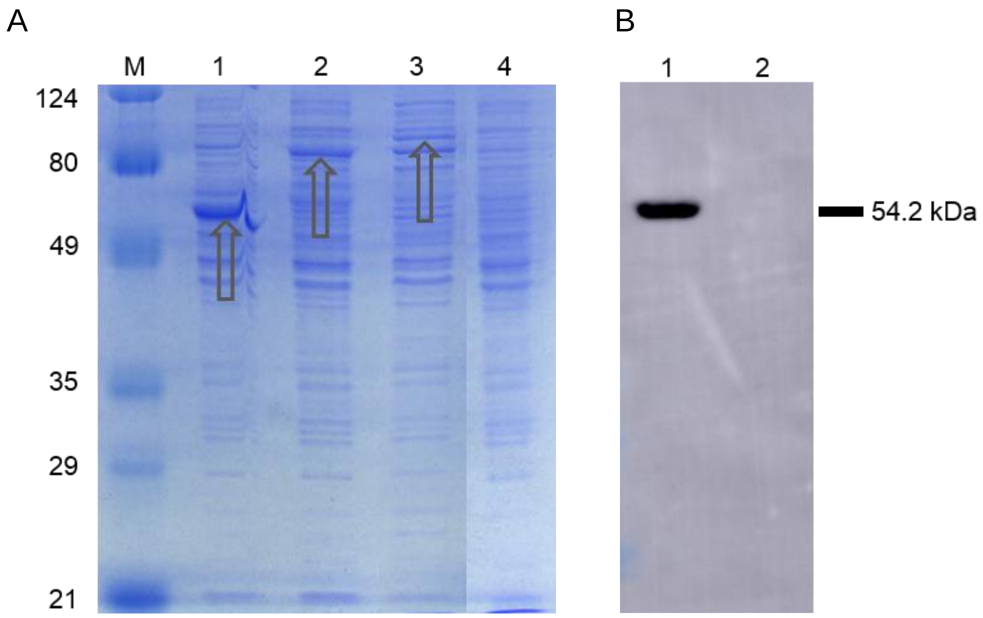


**Figure S1.** Expression analysis of β-glucosidases tested in *Pseudomonas putida* EM42. (**A**) Sodium dodecyl sulfate polyacrylamide gel electrophoresis (12 % gel) of cell free extracts obtained from *P. putida* EM42 pSEVA238_*bglC* (1), *P. putida* EM42 pSEVA238_*ccel2454* (2), *P. putida* EM42 pSEVA238_*bglX* (3), and host control *P. putida* EM42 pSEVA238 (4) grown in lysogeny broth and induced for 5 h with 1 mM 3-methylbenzoate. Theoretical molecular weights of β-glucosidases whose bands are signed with arrows are 53.4 kDa for BglC, 78.4 kDa for Ccel_2454, and 83.4 kDa for BglX. M is protein marker, values are in kDa. (**B**) Western blot analysis of BglC with N-terminal 6xHis tag in cell free extract obtained from *P. putida* EM42 pSEVA238_*bglC* (1) grown as described above, or in corresponding culture supernatant (2). Protein was detected using mouse anti-6xHis tag monoclonal antibody-HRP conjugate (Clontech, USA).

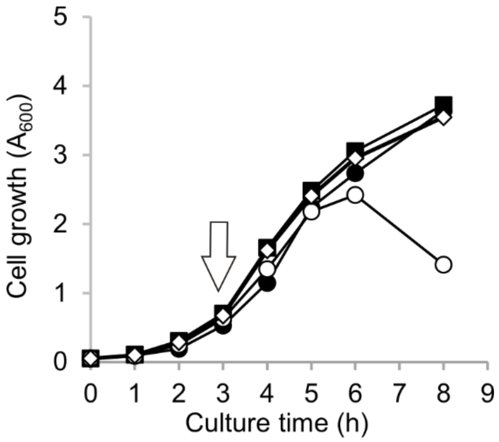


**Figure S2.** Effect of over-expression of tested β-glucosidases on growth of *P. putida* EM42 host. *P. putida* EM42 cells bearing pSEVA238 plasmid with one of three β-glucosidase genes were grown in lysogeny broth (30°C, 170 rpm). Expression was induced after 3 h of cultivation with 1 mM 3-methylbenzoate (marked by arrow) and cells were cultured for another 5 h in the same conditions. Empty pSEVA238 (control), filled squares (■); pSEVA238_*ccel2454*, filled circles (●); pSEVA238_*bglC*, open diamonds (◊); pSEVA238_*bglX*, open circles (○). Data points shown as means from two independent experiments.

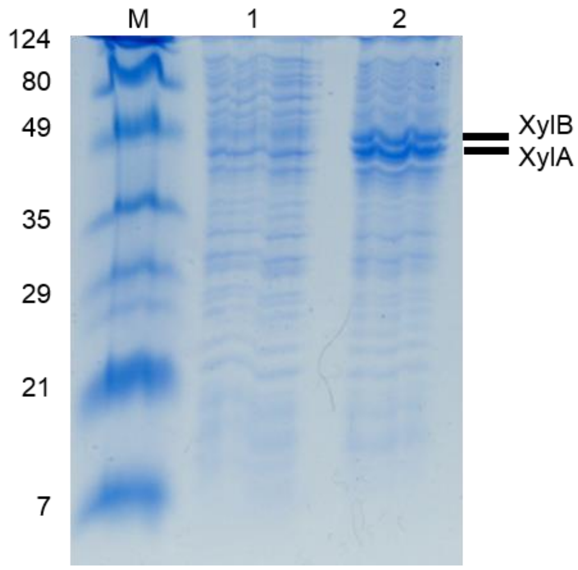


**Figure S3.** Sodium dodecyl sulfate polyacrylamide gel electrophoresis (12 % gel) of cell free extracts obtained from host control *P. putida* EM42 pSEVA2213 (1) or *P. putida* EM42 pSEVA2213_*xylAB* (2) grown overnight in lysogeny broth. Theoretical molecular weight of xylose isomerase XylA and xylulokinase XylB is 49.7 kDa and 52.6 kDa, respectively. M is protein marker, values are in kDa.

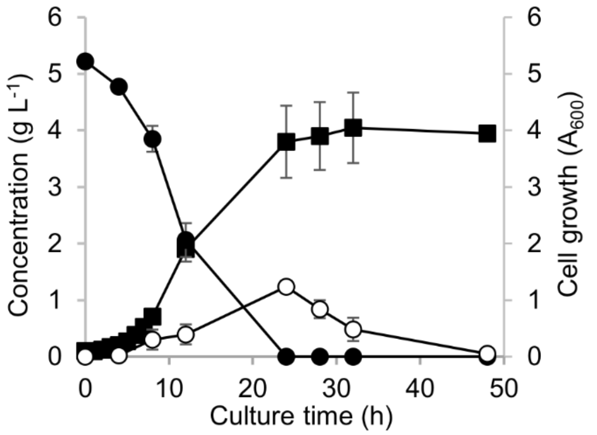


**Figure S4**. Growth of *P. putida* EM42 Δ*gcd bglC* in minimal medium with 5 g $L^{-1}$ D-cellobiose. Experiment was carried out in shaken flasks (30°C, 170 rpm). D-cellobiose, filled circles (●); D-glucose, open circles (○); cell growth, filled squares (■). Data points shown as mean ± SD from three independent experiments.

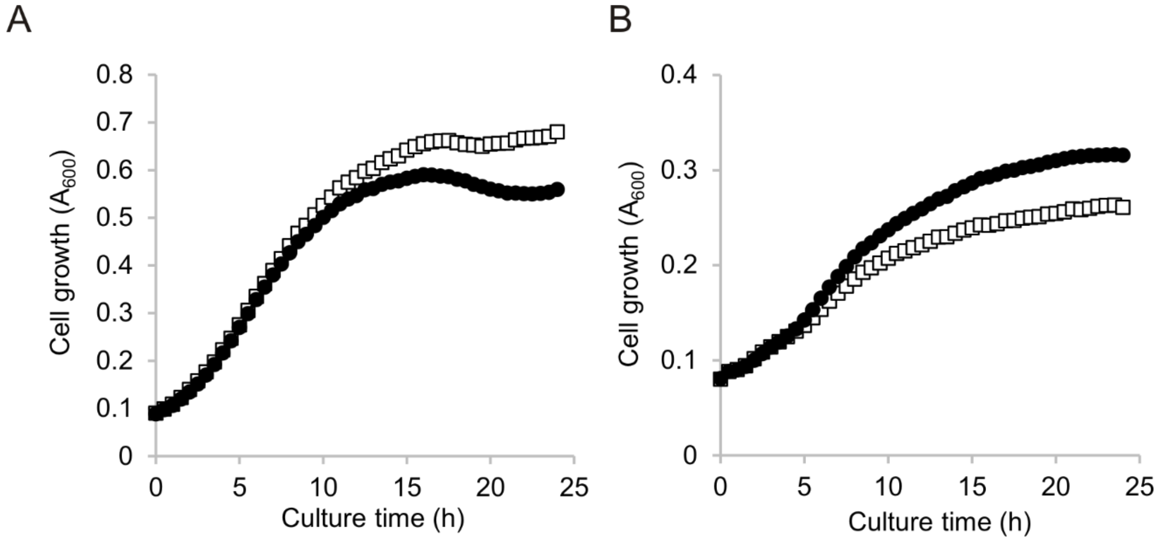


**Figure S5**. Comparison of growth of *P. putida* EM42 Δ*gcd* and *P. putida* EM42 Δ*gcd bglC* in rich growth medium and in minimal medium with gluconeogenic carbon source. (A) Rich lysogeny broth medium; (B) M9 minimal medium with citrate. Experiment was carried out in microtiter plate (150 μL of medium per well) at 30°C. *P. putida* EM42 Δ*gcd*, open squares (□); *P. putida* EM42 Δ*gcd bglC*, filled circles (●). Data points shown as means of absorbance $A_{600}$ measured at five wells. Standard deviations were within 10 % of the mean values.

**Table S1.** Oligonucleotide primers used in this study.

| | |
|---|---|
| bglX fw (NdeI) | TATACATATGATGAAACTGTCTTTGCTGG |
| bglX rv (HindIII) | ATTAAGCTTTTAAAGCAATTCAAAGCTCTGCTG |
| bglC fw (SacI) | AATGAGCTCGGTACCCTTTAAGAAG |
| bglC rv (PstI) | ATTCTGCAGCTATTCCTGTCCGAAGATTCC |
| xylAB fw (EcoRI) | TTCGAATTCTAGC*AAGAGG*AATATACC<br>ATGCAAGCCTATTTTGACCAG |
| xylAB rv (BamHI) | GATCGGATCCTTACGCCATTAATGGCAGAAG |
| xylE fw 1 | ATGAATACCCAGTATAATTCCAG |
| xylE rv 1 | TTACAGCGTAGCAGTTTGTTG |
| xylE fw 2 (BamHI) | AAAGGATCCCTTTAAG*AAGGAG*ATATACAT<br>ATGAATACCCAGTATAATTCC |
| xylE rv 2 (HindIII) | AAAAAGCTTTTACAGCGTAGCAGTTTG |
| gfpC fw (HindIII) | AATAAGCTTATGAGTAAAGGAGAAGAACTTTTCAC |
| gfpC rv (SpeI) | ATTACTAGTCTATTTGTATAGTTCATCCATGCC |
| xylE-gfp fw (BamHI) | ATTGGATCCCTTTAAGAAGGAGATATAC |
| xylE-gfp rv (HindIII) | ATTAAGCTTCAGCGTAGCAGTTTGTTGTG |
| Arb6 | GGCACGCGTCGACTAGTACNNNNNNNNNNACGCC |
| ME-O-Sm-Ext-F | CTTGGCCTCGCGCGCAGATCAG |
| Arb 2 | GGCACGCGTCGACTAGTAC |
| Me-O-Sm-Int-F | CACCAAGGTAGTCGGCAAAT |
| TS1F-gtsABCD (EcoRI) | GGAATTCGCATTGTTCGACACAGCCTG |
| TS1R-gtsABCD | TTATTGATGGTGTAGACGAGCGGAGCACCTTTCTTGTTGT |
| TS2F-gtsABCD | CTCGTCTACACCATCAATAA |
| TS2R-gtsABCD (BamHI) | CGGGATCCGTCGAAGTACTTCTGCTTGA |
| check(-)gtsABCD fw | GTGCCTTGCGTATCGCCGAA |
| gtsABCD junction check fw | GCTGCCGGATTACAACCTCG |
| gtsABCD junction check rv | GGCCAGTTGTACCAGATGCC |
| TS1F-gcd (EcoRI) | GGAATTCGCGGCAGTGCCGAGGTGTCGAAGTGGCGGTGG |
| TS1R-gcd | GGCCTGAAGATCCAGAGCAGTTTCTAACCCGCGACACCGCTCCC<br>GCAGGCTCAACCCTGAGG |
| TS2F-gcd | GGGTTAGAAACTGCTCTGGATCTTCAGGCC |
| TS2R-gcd (BamHI) | CGGGATCCGTCAGCCGGCCGCCCTCAGCGGCGCCGCCT |
| check(-)gcd fw | CTTCAGCTCTTCGCTGTACA |
| check(-)gcd rv | GCGTGCGCTACAACCTTAC |
| bglC check fw | CATCGTGCACGTGGACTAC |
| xerD check rv | CTGGATCAGGGCTACAAGC |

### Nucleotide sequences of genes used in this study

#### *ccel2_2454* (theoretical Mw of corresponding protein is 78.4 kDa)

ATGCAATACGATCAGATAGATAAAAAAATTGATGAACTGCTCTCTATGATGACTTTAGAGGAAAAGGCCGGCATGTGTCA
CGGAGCGGGACTTTTCAGAACCGCAGGAGTACCCAGATTGGGCATTCCTCCTCTGGTTTTTTCCGACGGCCCTATGGGGA
TCCGAAATGAGTTTGCTGACGATAATTGGAACACAGTGGGAGGAAATACAGATTTTGTTACATACCTTCCCGCCAATACCG
CACTTGCTGCAACCTTTAACCGGACTCTTGCTGAGAGTCTCGGAGAAGTACTGGGCTGCGAAGCACGAGGTCGGGGTAA
GGACGTCATCCTTGCTCCCGGAGTTAACATTATACGTACACCTTTATGCGGTAGAAATTATGAATACTTCAGTGAGGACCC
GATACTGACAGCTGAGCTTGCTGCTTCTTTTATAAAAGGTGTACAGCGTTTTGATGTGGCAGCCTGTGTCAAGCATTTTGC
TGCCAATAACCAGGAGACAGAACGTCTTGCCGTCAGTGCGGAGGTGGATGAACTTACACTGCGTGAATTGTACTTTCCCG
CATTTGAAGCATCAGTTCGGGCAGGAGTACTTACAGTTATGACTGCATATAACAGGCTCAATGGAACTTTTTGCAGCCATA
GCCGTCAGTTGATTACGGAGATCCTGCGTGAGGAATGGGGGTTTAACGGAGTTGTAGTATCGGACTGGGGTGCAGTACA
CGATACAGAATCCCCTGCAATTGCCGGACTTGACATTGAAATGAATGTAACCAGTAATTTTAATGAGTATTTCTTTGCAAA
ACCACTTATAAACGCAATCAAAGACGGAAAAATTCCTGAAAGGATGCTCGATGATAAGGTTCGTCGAATTCTGAGGCTGA
TGTTCAGACTTAACATGTTTTCAAAGGACCGTAAGCGTGGCGGCTTCAACTTACCGCAACACCAGCAGGCGGTTCTTGATG
CCGCTAAGGAAAGTTTTGTTCTTTTAAAAAATGACAGGGAAGTTCTGCCACTTAATGCGGACGGTATTAAAACAGTTGCTG
TTATCGGCAGTAATGCCGATAAAAAGCACTCTTCCGGGGGAGATAGTGCGGCAGTCAAGGCACTTTACGAGGTAACACC
GCTATCCGGCATAGTTATGAGGCTGGCAAGCGGAGCGAAGGTGACTTATTATCCGGGTTGTCCTGATGAAACACATTATA
AGGAGGAATTTCATATTCCGTCTAATGCTGATGAGAAAACACGTGCTGATATAGAGGAAAAGGCCAGAGCCGCTGATGA
AGACTACAGAAAAATCCAAATGCGACTTGAGGATGAGGCGATTCAGGCGGCTAAAACTGCCGATGCTGTTATTTTCATTG
GAGGGAACGGACATGAGCAGGAATCCGAAGGTCGTGATCGCCCGGATATGGCTTTGCCTTACGAACAGGATAAGCTGCT
GTCGCGTGTACTTGACGCAAATCCCAATACAGTAGTAGTTATTATAAGCGGCTCACCTGTTAATATGAGCGGCTGGATTGA
CAAGGCACCTACCGTAATGCAGGGATTTTTTTCTGGTATGCACGGCGGAACGGCTCTTGCGGCGGTTTTATTCGGAGACG
AAAATCCAAGCGGACATCTTCCCTTTACTATTCCCTTAAAAGAGGAGGAAACAGGAGCGAGTGCTCTGGGGGAATACCCG
GGCGGTGAAACGGTTTGTTACAGCGAGGGCTTGTTTGTGGGATACCGCTATCATGATGCATTCAACATACCTCCGCTTTTC
CCGTTTGGATATGGACTGTCGTATACCACTTTTTCATTGGCGAATGAATCCTTTAGGCGGCTACCTTGTGTCGGTACAGAG
TATGAAATCCATGTTGATATAACAAATTCCGGGAATAGGCCGGGTGCACAGTCAGTTCAGTTATATGTAGAGCCGGAAAA
AAAGGACGGCTCTCCGATACGTACTTTGAAAGGGTTTGAAAAGACATATCTCAATCCCGATGAAACCAAAACCATTACATT
TAAACTTGATGAACGTACATTCTCGGAATTCCGCCCCCATGAAGGCTGGGTTTTTGTACCGGGAAACTATACCATACATAT
TGGTACTTCGTCCCGTGAACTTCCGATAGCTATAGCTCTTGCTCTGTAA

#### *bglX* (theoretical Mw of corresponding protein is 83.4 kDa)

ATGATGAAACTGTCTTTGCTGGGCCTGGCCATGGGCCTTGCCAGTCAGGCGGCCCTCGCCGCCCCCACCGCCCCGCCCCTA
CAGGACAAGCAGGCCTTTATCGAGCATCTGATCAGCCAGATGACCGAAGCCGAAAAAATCGGCCAGTTGCGCCTGATCA
GCATCGGCCCGGAAATGCCCCGCGACAAGATCCGCGAGGAAATCGCCGCCGGCCGTATCGGTGGCACCTTCAACTCACG
CACTGCCCCCGAAAACCGGCCGATGCAGGACGCGGCCATGCGCAGCCGCCTGAAGATCCCGATGTTCTTCGCCTACGACA
CCGTCCACGGCGAGCGCACCATCTTCCCGATCGGCCTGGGCATGGCCGCGACCTGGGACATGGAGGCCGTCGCCAAGGT
CGGCCGCACCGCCGCGATCGAAGCCTCGGCCGACGCCCTGGACATGACGTTCGCGCCAATGGTGGATATCGCCCGTGAC
CCGCGCTGGGGCCGCACAAGCGAAGGTTTCGGCGAAGACACCTACCTGACTTCGAGAATCGGCCAGGTAATGGTGCGCT
CGTTCCAGGGCAGCAGCCCGGCCAACCCCGACAGCATCATGGCCATCGTCAAGCATTTCGCCTTGTATGGCGCCGTGGAA
GGCGGGCGCGACTACAACACGGTCGATATGAGCCTGCCGAAAATGTACAACGACTACCTGCCACCCTATCGCGCCGCGCT
CGATGCGGGTGCCGGTGGCGTGATGGTGGCGCTGAACTCGATCAACGGCGTACCGGCCACCTCCAACACCTGGCTGATG
AACGACCTGCTGCGCAAGGAGTGGGGCTTCAAGGGCGTGACCATCAGCGACCACGGTGCCATCCAGGAGCTGATTCGCC
ACGGCGTCGCCCGTGACGGCCGTGAAGCCGCCAAGCTGGCGATCAAGGCCGGCATCGACATGAGCATGAACGATACCCT
GTACGGCGAAGAGCTGCCGGGCCTGCTGAAGTCCGGCGAGGTGACCCAGCGCGAGCTGGACCAGGCGGTGCGTGAAGT
GCTGGGTGCCAAGTACGACATGGGCCTGTTCAAGGACCCGTACGTGCGCATCGGCAAGGCCGAAACCGATCTGAAGGAC
TACTACGGCAACGACCGCCTGCACCGCGAAGCTGCGCGCGACGTGGCACGCCGCAGCCTGGTGCTGCTGGAAAACCGCA
ACCAGACCCTGCCGCTGAAAAAGGCCGGCACCATTGCCTTGGTCGGCCCGCTGGCCGATGCTCCGATCGACATGATGGGC
AGCTGGGCCGCCGACGGTAAACCCGTCCACTCGGTGACCGTGCGCGAAGGCTTGCGCCGCGCCGTGGAAGGCAAGGCC
AAGCTGGTCTACGCCAAAGGCTCCAACGTCACGGGCGACAAGGCGATCTTCGATTACCTGAACTTCCTCAACTTCGATGCC
CCGGAAATCGTCGACGACCCACGCCCGCCTGCGGTGCTGATCGATGAAGCAATCAAGGCAGCCAAGCAATCCGACGTCG
TGGTCGCAGTGGTCGGCGAGTCCCGCGGCATGTCCCACGAGTCGTCGAGCCGCACCACCCTGGAGATCCCGGCCAGCCA

GCGCGAGCTGATCAAGGCGCTCAAGGCCACCGGCAAACCGCTGGTGCTGGTGCTGATGAACGGCCGGCCGCTGTCGATC
AGCTGGGAGCGCGAGCAGGCCGACGCCATCCTGGAAACCTGGTTCGCCGGCACCGAAGGCGGCAATGCCATCGCCGAC
GTGCTGTTTGGCGACTACAACCCCTCCGGCAAGCTGGCCATTACCTTCCCACGGTCGGTGGGGCAGATCCCGATGTACTA
CAACCACACCCGCATCGGCCGGCCGTTCACGCCGGGCAAGCCGGGCAACTACACCTCGCAATACTTCGAAGAACCCAACG
GGCCGCTTTATCCGTTCGGCTATGGCTTGAGCTACAGCAGCTTCGAGCTGTCAGGCCTGAACCTGTCGAGCAAGGATCTC
AAGCGCGGCGATACGCTGGATGCCAAGGTAACGGTCAAGAACACCGGCAAGGTTGCAGGCGAGACGGTGGTGCAGCTG
TACCTGCAGGATGTGTCGGCGTCGATGAGCCGCCCAGTGAAAGAGCTGAAAAACTTCCAGAAGCTGATGCTTGAACCGG
GTGAAACCCGCACGTTGACCTTCCGCATCAGCGAGGACGACCTGAAGTTCTACAATGGCCAGTTGCAGCGGGTTGCCGAA
CCAGGGGAGTTCAATGTCCAGGTAGGGCTGGATTCCCAGGCGGTGCAGCAGCAGAGCTTTGAATTGCTTTAA

***bglC* (theoretical Mw of corresponding protein is 53.4 kDa)**

ATGACCTCGCAATCGACGACTCCTCTGGGCAATCTCGAGGAGACTCCCAAACCGGATATCCGCTTCCCGTCCGATTTCGTG
TGGGGAGTGGCGACCGCTTCGTTCCAGATCGAAGGCTCCACCACGGCCGACGGCCGCGGCCCCAGCATCTGGGACACCT
TCTGCGCCACTCCGGGCAAGGTCGAGAACGGCGACACGGGCGACCCTGCCTGCGACCACTACAACCGGTACCGCGATGA
CGTGGCCTTGATGCGGGAGCTGGGCGTGGGCGCCTACCGCTTCTCCATCGCCTGGCCGCGGATCCAGCCCGAGGGCAAG
GGCACGCCCGTGGAGGCCGGGCTGGACTTCTACGACCGGCTTGTGGACTGCCTGCTGGAGGCCGGCATCGAGCCGTGGC
CGACCCTCTACCACTGGGACCTGCCGCAGGCGCTGGAGGACGCGGGCGGCTGGCCCAACCGGGACACGGCCAAGCGGT
TCGCCGACTACGCGGAGATCGTCTACCGCCGGCTCGGCGACCGGATCACCAACTGGAACACGCTCAACGAGCCGTGGTG
CTCCGCGTTCCTGGGCTACGCCTCCGGCGTGCACGCCCCGGGCCGCCAGGAGCCGGCTGCTGCGCTGGCCGCCGCCCACC
ACCTGATGCTGGGCCACGGGCTGGCCGCTGCCGTGATGCGGGACTTGGCGGGCCAGGCCGGACGTTCCGTGCGGATCG
GTGTCGCGCACAACCAGACCACGGTCCGTCCCTACACTGACAGTGAGGCCGACCGGGACGCTGCGCGCCGGATTGACGC
CCTGCGGAACCGCATCTTCACCGAGCCGCTGGTGAAGGGCCGCTACCCGGAGGACCTGATCGAGGACGTCGCCGCGGTC
ACCGACTACAGCTTCGTCCAGGACGGCGACCTGAAGACCATCTCCGCCAACCTGGACATGATGGGCGTCAACTTCTACAA
CCCGAGCTGGGTGTCAGGCAACCGGGAGAACGGGGGCTCCGACCGGCTGCCCGACGAGGGCTACTCGCCGTCGGTCGG
CAGCGAGCATGTCGTGGAGGTGGACCCCGGCCTGCCGGTGACCGCCATGGGCTGGCCGATCGACCCGACCGGGCTGTAC
GACACGCTGACCCGGCTGGCCAACGACTACCCGGGCCTGCCGCTGTACATCACCGAGAACGGCGCCGCCTTCGAGGACA
AGGTGGTCGACGGCGCGGTGCACGACACCGAGCGGATCGCCTACCTGGACTCGCACCTGCGGGCCGCGCACGCTGCCAT
TGAGGCGGGCGTGCCGCTCAAGGGCTACTTCGCCTGGTCGTTCATGGACAACTTCGAGTGGGCCCTCGGGTACGGGAAG
CGGTTCGGCATCGTGCACGTGGACTACGAGAGCCAGACGCGCACGGTGAAGGACAGCGGCTGGTGGTACTCCCGGGTG
ATGCGCAACGGGGGAATCTTCGGACAGGAATAG

***xylA* (theoretical Mw of corresponding protein is 49.7 kDa)**

ATGCAAGCCTATTTTGACCAGCTCGATCGCGTTCGTTATGAAGGCTCAAAATCCTCAAACCCGTTAGCATTCCGTCACTACA
ATCCCGACGAACTGGTGTTGGGTAAGCGTATGGAAGAGCACTTGCGTTTTGCCGCCTGCTACTGGCACACCTTCTGCTGG
AACGGGGCGGATATGTTTGGTGTGGGGGCGTTTAATCGTCCGTGGCAGCAGCCTGGTGAGGCACTGGCGTTGGCGAAG
CGTAAAGCAGATGTCGCATTTGAGTTTTTCCACAAGTTACATGTGCCATTTTATTGCTTCCACGATGTGGATGTTTCCCCTG
AGGGCGCGTCGTTAAAAGAGTACATCAATAATTTTGCGCAAATGGTTGATGTCCTGGCAGGCAAGCAAGAAGAGAGCGG
CGTGAAGCTGCTGTGGGGAACCGCCAACTGCTTTACAAACCCTCGCTACGGCGCGGGTGCGGCGACGAACCCAGATCCT
GAAGTCTTCAGCTGGGCGGCAACGCAAGTTGTTACAGCGATGGAAGCAACCCATAAATTGGGCGGTGAAAACTATGTCC
TGTGGGGCGGTCGTGAAGGTTACGAAACGCTGTTAAATACCGACTTGCGTCAGGAGCGTGAACAACTGGGCCGCTTTAT
GCAGATGGTGGTTGAGCATAAACATAAAATCGGTTTCCAGGGCACGTTGCTTATCGAACCGAAACCGCAAGAACCGACCA
AACATCAATATGATTACGATGCCGCGACGGTCTATGGCTTCCTGAAACAGTTTGGTCTGGAAAAAGAGATTAAACTGAAC
ATTGAAGCTAACCACGCGACGCTGGCAGGTCACTCTTTCCATCATGAAATAGCCACCGCCATTGCGCTTGGCCTGTTCGGT
TCTGTCGACGCCAACCGTGGCGATGCGCAACTGGGCTGGGACACCGACCAGTTCCCGAACAGTGTGGAAGAGAATGCGC
TGGTGATGTATGAAATTCTCAAAGCAGGCGGTTTCACCACCGGTGGTCTGAACTTCGATGCCAAAGTACGTCGTCAAAGT
ACTGATAAATATGATCTGTTTTACGGTCATATCGGCGCGATGGATACGATGGCACTGGCGCTGAAAATTGCAGCGCGCAT
GATTGAAGATGGCGAGCTGGATAAACGCATCGCGCAGCGTTATTCCGGCTGGAATAGCGAATTGGGCCAGCAAATCCTG
AAAGGCCAAATGTCACTGGCAGATTTAGCCAAATATGCTCAGGAACATAATTTGTCTCCGGTGCATCAGAGTGGTCGCCA
GGAGCAACTGGAAAATCTGGTAAATCATTATCTGTTCGACAAATAA

***xylB* (theoretical Mw of corresponding protein is 52.6 kDa)**

ATGTATATCGGGATAGATCTTGGCACCTCGGGCGTAAAAGTTATTTTGCTCAACGAGCAGGGTGAGGTGGTTGCTTCGCA
AACGGAAAAGCTGACCGTTTCGCGCCCGCATCCACTCTGGTCGGAACAAGACCCGGAACAGTGGTGGCAGGCAACTGAT
CGCGCAATGAAAGCTCTGGGCGATCAGCATTCTCTGCAGGACGTTAAAGCATTGGGTATTGCCGGCCAGATGCATGGAG
CAACCTTACTGGATGCTCAACAACGGGTATTGCGCCCTGCCATTTTGTGGAACGACGGGCGCTGTGCGCAAGAGTGCACT
TTGCTGGAAGCGAGAGTTCCGCAATCACGAGTGATTACCGGCAACCTGATGATGCCCGGATTTACTGCGCCTAAATTGCT
ATGGGTTCAGCGGCATGAGCCGGAGATATTCCGTCAAATCGACAAAGTATTATTACCGAAAGATTACTTGCGTCTGCGTA
TGACGGGGGAGTTTGCCAGCGATATGTCTGACGCAGCTGGCACCATGTGGCTGGATGTCGCAAAGCGTGACTGGAGTGA
CGTCATGCTGCAGGCTTGCGACTTATCTCGTGACCAGATGCCCGCATTATACGAAGGCAGCGAAATTACTGGTGCTTTGTT
ACCTGAAGTTGCGAAAGCGTGGGGTATGGCGACGGTGCCAGTTGTCGCAGGCGGTGGCGACAATGCAGCTGGTGCAGT
TGGTGTGGGAATGGTTGATGCTAATCAGGCAATGTTATCGCTGGGGACGTCGGGGGTCTATTTTGCTGTCAGCGAAGGG
TTCTTAAGCAAGCCAGAAAGCGCCGTACATAGCTTTTGCCATGCGCTACCGCAACGTTGGCATTTAATGTCTGTGATGCTG
AGTGCAGCGTCGTGTCTGGATTGGGCCGCGAAATTAACCGGCCTGAGCAATGTCCCAGCTTTAATCGCTGCAGCTCAACA
GGCTGATGAAAGTGCCGAGCCAGTTTGGTTTCTGCCTTATCTTTCCGGCGAGCGTACGCCACACAATAATCCCCAGGCGA
AGGGGGTTTTCTTTGGTTTGACTCATCAACATGGCCCCAATGAACTGGCGCGAGCAGTGCTGGAAGGCGTGGGTTATGCG
CTGGCAGATGGCATGGATGTCGTGCATGCCTGCGGTATTAAACCGCAAAGTGTTACGTTGATTGGGGGCGGGGCGCGTA
GTGAGTACTGGCGTCAGATGCTGGCGGATATCAGCGGTCAGCAGCTCGATTACCGTACGGGAGGGGATGTGGGGCCAG
CACTGGGCGCAGCAAGGCTGGCGCAGATCGCGGCGAATCCAGAGAAATCGCTCATTGAATTGTTGCCGCAACTACCGTTA
GAACAGTCGCATCTACCAGATGCGCAGCGTTATGCCGCTTATCAGCCACGACGAGAAACGTTCCGTCGCCTCTATCAGCA
ACTTCTGCCATTAATGGCGTAA

***xylE* (theoretical Mw of corresponding protein is 53.6 kDa)**

ATGAATACCCAGTATAATTCCAGTTATATATTTTCGATTACCTTAGTCGCTACATTAGGTGGTTTATTATTTGGCTACGACAC
CGCCGTTATTTCCGGTACTGTTGAGTCACTCAATACCGTCTTTGTTGCTCCACAAAACTTAAGTGAATCCGCTGCCAACTCC
CTGTTAGGGTTTTGCGTGGCCAGCGCTCTGATTGGTTGCATCATCGGCGGTGCCCTCGGTGGTTATTGCAGTAACCGCTTC
GGTCGTCGTGATTCACTTAAGATTGCTGCTGTCCTGTTTTTTATTTCTGGTGTAGGTTCTGCCTGGCCAGAACTTGGTTTTA
CCTCTATAAACCCGGACAACACAGTGCCTGTTTATCTGGCAGGTTATGTCCCGGAATTTGTTATTTATCGCATTATTGGCGG
TATTGGCGTTGGTTTAGCCTCAATGCTCTCGCCAATGTATATTGCGGAACTGGCTCCAGCTCATATTCGCGGGAAACTGGT
CTCTTTTAACCAGTTTGCGATTATTTTCGGGCAACTTTTAGTTTACTGCGTAAACTATTTTATTGCCCGTTCCGGTGATGCCA
GCTGGCTGAATACTGACGGCTGGCGTTATATGTTTGCCTCGGAATGTATCCCTGCACTGCTGTTCTTAATGCTGCTGTATAC
CGTGCCAGAAAGTCCTCGCTGGCTGATGTCGCGCGGCAAGCAAGAACAGGCGGAAGGTATCCTGCGCAAAATTATGGGC
AACACGCTTGCAACTCAGGCAGTACAGGAAATTAAACACTCCCTGGATCATGGCCGCAAAACCGGTGGTCGTCTGCTGAT
GTTTGGCGTGGGCGTGATTGTAATCGGCGTAATGCTCTCCATCTTCCAGCAATTTGTCGGCATCAATGTGGTGCTGTACTA
CGCGCCGGAAGTGTTCAAAACGCTGGGGGCCAGCACGGATATCGCGCTGTTGCAGACCATTATTGTCGGAGTTATCAACC
TCACCTTCACCGTTCTGGCAATTATGACGGTGGATAAATTTGGTCGTAAGCCACTGCAAATTATCGGCGCACTCGGAATGG
CAATCGGTATGTTTAGCCTCGGTACCGCGTTTTACACTCAGGCACCGGGTATTGTGGCGCTACTGTCGATGCTGTTCTATG
TTGCCGCCTTTGCCATGTCCTGGGGTCCGGTATGCTGGGTACTGCTGTCGGAAATCTTCCCGAATGCTATTCGTGGTAAAG
CGCTGGCAATCGCGGTGGCGGCCCAGTGGCTGGCGAACTACTTCGTCTCCTGGACCTTCCCGATGATGGACAAAAACTCC
TGGCTGGTGGCCCATTTCCACAACGGTTTCTCCTACTGGATTTACGGTTGTATGGGCGTTCTGGCAGCACTGTTTATGTGG
AAATTTGTCCCGGAAACCAAAGGTAAAACCCTTGAGGAGCTGGAAGCGCTCTGGGAACCGGAAACGAAGAAAACACAAC
AAACTGCTACGCTGTAA
